# Surface-active antibiotic production is a multifunctional adaptation for postfire microbes

**DOI:** 10.1101/2023.08.17.553728

**Authors:** Mira D. Liu, Yongle Du, Sara K. Koupaei, Nicole R. Kim, Wenjun Zhang, Matthew F. Traxler

**Author notes:** Corresponding author: Matthew F. Traxler **Email:**. **Author contributions:** Mira D. Liu: Conceptualization, Investigation, Data Curation, Funding Acquisition, Formal Analysis, Supervision, Writing – original draft, Writing – review & editing Yongle Du: Formal Analysis, Writing – original draft Sara K. Koupaei: Investigation Nicole R. Kim: Investigation Wenjun Zhang: Writing – review & editing Matthew F. Traxler: Conceptualization, Supervision, Funding Acquisition, Project administration, Writing – original draft, Writing – review & editing. **Competing interest statement:** The authors declare no competing interests. **Classification:** Major: Biological Sciences, Minor: Microbiology.

## Abstract

Wildfires affect soils in multiple ways, leading to numerous challenges for colonizing microbes. While it is thought that fire-adapted microbes lie at the forefront of postfire ecosystem recovery, the specific strategies that these microbes use to thrive in burned soils remain largely unknown. Through bioactivity screening of bacterial isolates from burned soils, we discovered that several *Paraburkholderia spp.* isolates produced a set of unusual rhamnolipid surfactants with a natural methyl ester modification. These rhamnolipid methyl esters (RLMEs) exhibited enhanced antimicrobial activity against other postfire microbial isolates, including pyrophilous *Pyronema* fungi and *Amycolatopsis* bacteria, compared to the typical rhamnolipids made by organisms such as *Pseudomonas spp*. RLMEs also showed enhanced surfactant properties and facilitated bacterial motility on agar surfaces. *In vitro* assays further demonstrated that RLMEs improved aqueous solubilization of polycyclic aromatic hydrocarbons, which are potential carbon sources found in char. Identification of the rhamnolipid biosynthesis genes in the postfire isolate, *Paraburkholderia caledonica* str. F3, led to the discovery of *rhlM*, whose gene product is responsible for the unique methylation of rhamnolipid substrates. RhlM is the first characterized bacterial representative of a large class of integral membrane methyltransferases that are widespread in bacteria. These results indicate multiple roles for RLMEs in the postfire lifestyle of *Paraburkholderia* isolates, including enhanced dispersal, solubilization of potential nutrients, and inhibition of competitors. Our findings shed new light on the chemical adaptations that bacteria employ in order to navigate, grow, and outcompete other soil community members in postfire environments.

**Significance Statement:** Wildfires are increasing in frequency and intensity at a global scale. Microbes are the first colonizers of soil after fire events, but the adaptations that help these organisms survive in postfire environments are poorly understood. In this work, we show that a bacterium isolated from burned soil produces an unusual rhamnolipid biosurfactant that exhibits antimicrobial activity, enhances motility, and solubilizes potential nutrients derived from pyrolyzed organic matter. Collectively, our findings demonstrate that bacteria leverage specialized metabolites with multiple functions to meet the demands of life in postfire environments. Furthermore, this work reveals the potential of probing perturbed environments for the discovery of unique compounds and enzymes.

## Introduction

Wildfires represent an archetypal disturbance regime that affects communities of animals, plants, and microorganisms. The below-ground effects of fire on the soil nutrient landscape are stratified, with the most intense alterations occurring at the surface. In the top layer of soil, fire reduces the amount of bioavailable carbon as extreme heat leads to combustive release of carbon dioxide, and most of the remaining carbon is converted into pyrolyzed organic matter (PyOM)(1, 2). PyOM predominantly consists of stable, aromatic C products including polycyclic aromatic hydrocarbons (PAHs). These recalcitrant molecules are not only difficult to degrade, but are also hydrophobic and poorly bioavailable.

Organisms from fire-prone ecosystems often possess adaptations that enable survival or rapid recolonization after fire. For example, many conifers produce thick protective bark and fire-activated serotinous cones (3). Previous research suggests that soil microbes lie at the forefront of postfire community recovery processes (4). Unlike most plants and animals, some bacteria and fungi can access pyrolyzed chemical products and reintegrate them into the local food web, as in the case of fungi such as *Pyronema spp* (5, 6). However, microbes face numerous other challenges in a burned soil environment beyond nutrient limitation, such as poor nutrient solubility, increased hydrophobicity of the local environment, and competition with other fire-adapted organisms. Thus, PyOM-adapted catabolism alone may be an insufficient strategy to thrive given the multifaceted demands presented by the postfire environment.

After a perturbation such as fire, organismal communities can recover over time through the process of ecological succession. In perturbed systems with limited or altered nutrient availability, competitive interactions between colonizing members may play an important role in shaping the developing community. Early microbial colonizers, such as certain species of fungi, are termed “pyrophilous” because they consistently emerge after fire. The fungal genus *Pyronema* contains multiple pyrophilous species that often rapidly dominate postfire microbial communities (7). However, after a short-lived peak, these *Pyronema* species sharply decline in abundance. Numerous factors may be responsible for the decline of *Pyronema*, such as nutrient depletion or changes in pH. Alternatively, the rapid decline of *Pyronema* might result from competitive interspecies interactions.

In this work, we sought to explore the possibility that interference competition mediated by specialized metabolism, *i.e.* production of inhibitory secondary metabolites, could play a role in postfire microbial interactions. We also considered whether such specialized metabolites might confer other adaptive advantages within postfire environments. Recent work has underscored the notion that specialized metabolites may have multiple functions in nutrient limited environments. For instance, microbially produced phenazines have long been recognized as antimicrobial agents, but recently have been shown to play roles in enhancing the bioavailability of phosphorus (8) and iron (9), and in redox balancing under anaerobic conditions (10). Thus, we hypothesized that adaptations of postfire microorganisms may present new opportunities for 1) discovery of novel specialized metabolites, and 2) understanding the roles of specialized metabolites *in situ*.

To explore these possibilities, we screened for molecules produced by bacteria isolated from postfire soils that could inhibit the growth of *Pyronema*. We report that multiple postfire isolates of the genus *Paraburkholderia* produced novel members of a class of biosurfactants, rhamnolipid methyl esters (RLMEs), which likely provide multiple adaptive advantages for their producers. Collectively, our findings shed light on chemical adaptations that bacteria may employ in burned soil environments in order to grow, survive, and outcompete other community members. Beyond this, our results highlight microbes from perturbed environments as fruitful sources for discovery of novel compounds and enzymes.

## Results

### Screening burned soil bacterial isolates for activity against *Pyronema omphalodes*

The pyrophilous fungus *Pyronema omphalodes* has been observed to dominate the postfire fungal community, but then rapidly decline in abundance (7). We hypothesized that specialized metabolite-mediated antagonism may be one factor contributing to *Pyronema*’s decline. Therefore, we screened a library of bacteria isolated from soil collected from Blodgett Forest Research Station, which underwent a prescribed burn in October 2018, for antifungal activity against *P. omphalodes* using an agar plug assay (Fig. 1A). To mimic environmental conditions, we grew the bacterial isolates on solid minimal medium containing PyOM as the sole carbon source. We also grew each strain in parallel on a rich medium (ISP2 agar), which contained glucose as the carbon source. For each isolate screened, we looked for differential antifungal activity from plugs that were taken from PyOM cultures compared to ISP2 cultures. We hypothesized that isolates exhibiting increased bioactivity when grown on PyOM were more likely to be sources of novel compounds, which may have remained undetected using traditional, rich medium-based laboratory screens.

**Figure 1.**
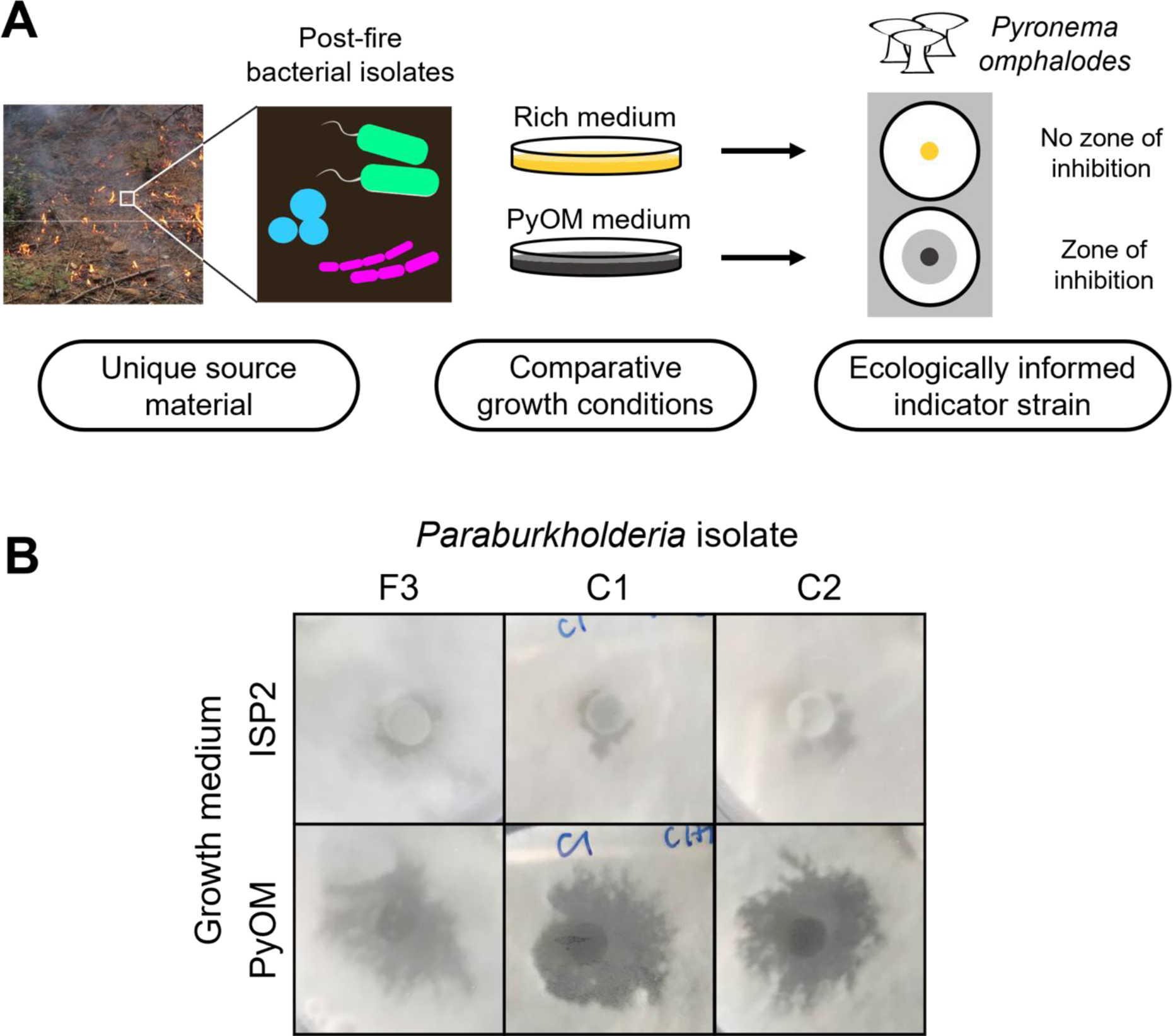
(A) Ecologically based natural product discovery framework. Bacterial isolates from postfire soils were cultivated in parallel on a rich medium and PyOM-containing medium. Solid cultures were screened using an agar plug assay for antifungal activity against *Pyronema omphalodes*. (B) Preliminary antifungal screening results for *Paraburkholderia* isolates F3, C1, and C2 cultivated on PyOM agar and a rich medium (International Streptomyces Project 2, ISP2 agar), assayed against *Pyronema omphalodes* P1672. Zones of inhibition from *Paraburkholderia*-produced metabolites were larger from PyOM cultures than ISP2 cultures.

Out of 38 bacterial isolates screened, 30 strains displayed antifungal activity against *Pyronema omphalodes.* Five strains produced larger zones of inhibition when grown on PyOM (results for strains F3, C1, C2 shown in Fig.1B), all of which were identified as members of the genus *Paraburkholderia* by amplification of their 16S rRNA genes. The results from this screen revealed that antifungal activity is prevalent among bacterial postfire soil isolates, and *Paraburkholderia* isolates are particularly noteworthy in their heightened activity during culturing on PyOM agar.

### Discovery and structural elucidation of novel antibiotic surfactants: rhamnolipid methyl esters

Based on the above antifungal screen, we selected isolate F3, tentatively identified here as *Paraburkholderia caledonica*, as a promising candidate for further chemical analysis and investigation, with the aim to identify the molecule responsible for *P. omphalodes* inhibition. We scaled-up cultivation of *P. caledonica* F3, extracted the spent medium with ethyl acetate, and subjected the crude extract to preliminary purification *via* solid-phase extraction (SPE).

Subsequent antifungal assay-guided fractionation *via* semi preparative-scale reverse-phase high pressure liquid chromatography (RP-HPLC) led to the isolation of the active molecule whose structure is shown in Fig. 2A. Positive-mode high-resolution mass spectrometry (HRMS) analysis revealed a peak for an ammonium adduct ion at *m/z* 738.4995, corresponding to a neutral species molecular formula of C_37_H_68_O_13_. The compound was proposed as a rhamnolipid through 1D and 2D nuclear magnetic resonance (NMR) spectroscopy and comparison with a previous reference (Table S4, Fig. S1-4) (11). The presence of the methyl ester was indicated by ^3^*J*-HMBC correlation from H-11 (*δ*_H_ 3.58) to C-1 (*δ*_C_ 170.4) (Fig. S4). The length of the two lipid chains, presence of rhamnose groups, stereochemistry, and connections between these moieties were further elucidated as detailed in the Materials and Methods (Table S4, Fig. S1-4). High resolution tandem mass spectrometry (HR-MS/MS) analysis enabled the further identification of additional RLME analogs, with varying lengths in the second acyl chain (Fig. 2C). These analogs were named RLME A-C according to decreasing observed relative abundances. Together, these data indicated that the active molecules (1–3) were rhamnolipids carrying an unusual methyl ester modification where typical rhamnolipids, such as Rhamnolipid 1 produced by *Pseudomonas sp.* (4), terminate in a carboxyl group. This methylation, which alters the polarity of the molecule, is expected to impart altered surfactant properties to these RLMEs.

**Figure 2.**
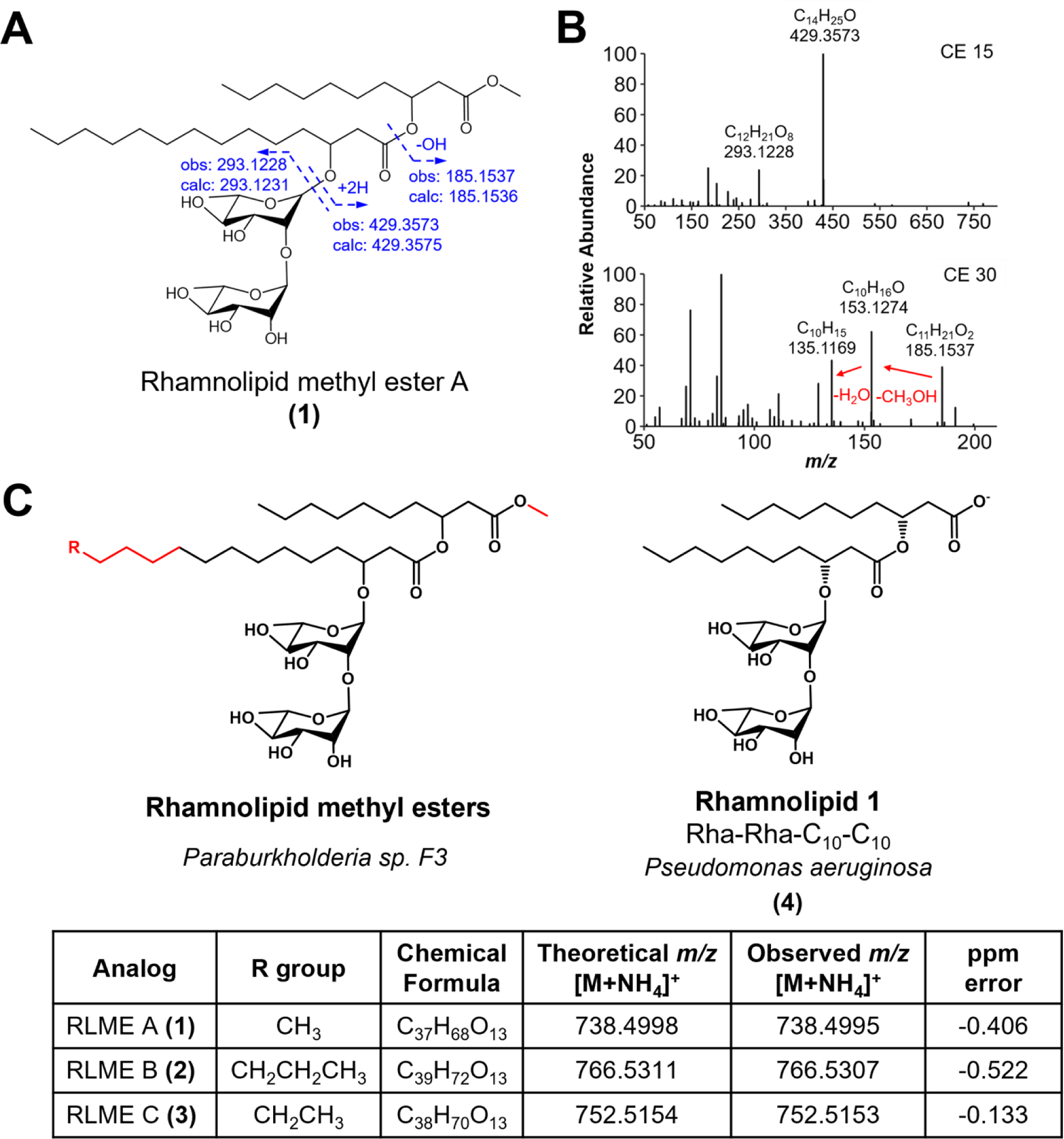
Structural elucidation of novel rhamnolipid methyl esters. **(A)** Proposed chemical structure for the *Paraburkholderia caledonia* str. F3-produced rhamnolipid methyl ester A. Dashed arrows indicate theoretical molecular ion fragments that produce *m/z* values observed (obs) in the experimental data shown in (B) along with theoretically calculated values (cal). **(B)** HR-MS/MS fragmentation spectra obtained using collision energy of 15 eV (top) and 30 eV (bottom). **(C)** Rhamnolipid methyl esters produced by *P. caledonica* F3 with details for each analog, and the known Rhamnolipid 1 produced by *Pseudomonas aeruginosa*. Structural differences are highlighted in red. Theoretical *m/z* values and ppm errors were calculated using the Barrow group online calculator tool. Chemical formulas represent neutral species.

### Identification of a novel rhamnolipid methyltransferase encoded by *rhlM*

In order to identify the genes responsible for RLME biosynthesis, and in particular rhamnolipid carboxyl methylation, we sequenced the full genome of *Paraburkholderia caledonica* F3. A BLAST search of the *P. caledonica* F3 genome using *Pseudomonas aeruginosa rhl* gene homologs led us to identify the *rhl* gene cluster (Fig. 3A-B). The uncharacterized gene downstream of *rhlA* was annotated as an isoprenylcysteine carboxyl methyltransferase family protein. We hypothesized that the enzyme encoded by this gene likely catalyzed the methylation of rhamnolipids to form rhamnolipid methyl esters.

**Figure 3.**
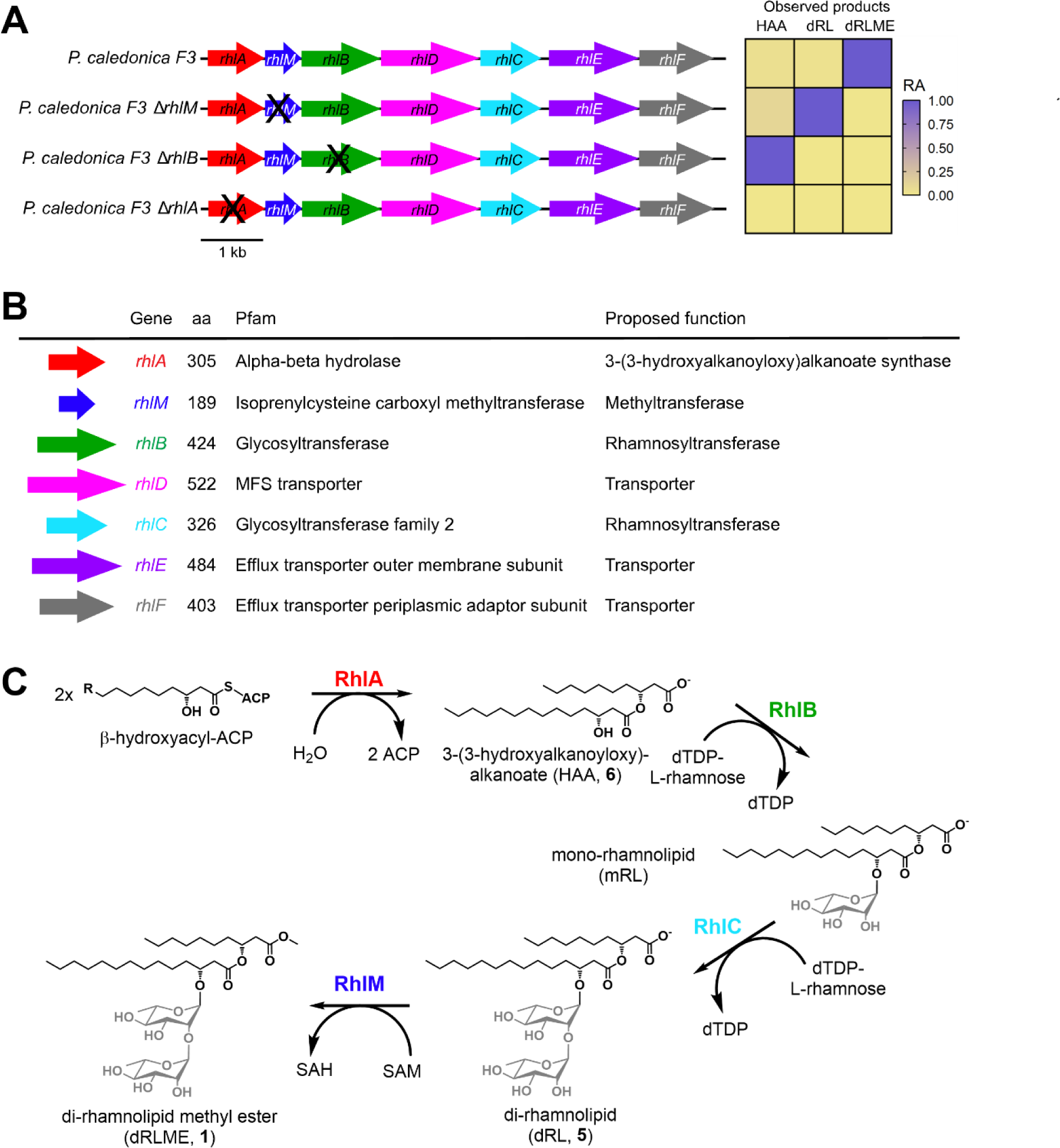
Rhamnolipid methyl ester biosynthesis genes are clustered in *Paraburkholderia caledonica* F3. (**A)** Genotypes of WT and knockout mutants. Heatmap of production of RLME and biosynthetic intermediates of *P. caledonica* F3 wild type and knockout mutants, observed using LC-MS. Relative abundance (RA) of possible *rhl* pathway intermediates found in *P. caledonica* F3 knockout strains and wild type (WT), by extracted ion chromatogram peak area and normalized by relative abundance (RA) within each strain (slate blue, more abundant; tan, less abundant). Genes in the RLME biosynthetic gene cluster with their amino acid (aa) length, Pfam annotation, and proposed function. The newly identified *rhlM* gene encodes for a putative Class VI integral membrane methyltransferase. **(C)** Proposed pathway for biosynthesis of di-rhamnolipid methyl ester A in *P. caledonica* F3. HAA, 3-(3-hydroxyalkanoyloxy)alkanoate; dRL, di-rhamnolipid; dRLME, dirhamnolipid methyl ester; ACP, acyl carrier protein; dTDP, deoxythymidine diphosphate; mRL, mono-rhamnolipid.

To test this hypothesis, we used double allelic exchange to create a knockout mutant lacking the *rhlM* gene. Furthermore, to guide the full elucidation of the biosynthetic pathway, we created single mutants of *rhlA* and *rhlB* for chemical analysis of their intermediate products. We grew each mutant alongside wildtype *P. caledonica* F3 (WT), performed chemical extractions using ethyl acetate, and analyzed the extracts using liquid chromatography coupled with tandem mass spectrometry (LC-MS/MS).

Deletion of *rhlM* abolished production of rhamnolipid methyl esters (Fig. 3A), while precursors including O-desmethyl rhamnolipid products and 3-(3-hydroxyalkanoyloxy)alkanoates (HAAs) were still produced (Fig. S5). Complementation of Δ*rhlM* with a *rhlM*-expressing plasmid rescued production of RLMEs, confirming the role of *rhlM* in methylation of rhamnolipid precursors. As predicted, in Δ*rhlB* strain extracts, HAAs were detected while rhamnosylated products were not (Fig. 3A, Fig. S5). No rhamnolipid pathway intermediates were detected in Δ*rhlA* extracts (Fig. 3A, Fig. S5). These results confirmed the roles of *rhlA* and *rhlB* in 3-hydroxy-fatty acid esterification and rhamnosylation, respectively.

These phenotypes were similarly rescued with their respective complementation on stable plasmids with constitutive expression (pBBR1-MCS5) (Fig. S5). Together, the observed intermediate products of Δ*rhlA,* Δ*rhlB,* and Δ*rhlM* allow us to propose rhamnolipid methylation by RhlM as the final step in the biosynthesis of RLMEs (Fig. 3C). Furthermore, in order to determine whether *rhlM*-dependent RLME production is strain-specific or is widespread in the *Paraburkholderia* clade, we examined 6 additional *Paraburkholderia* isolates from burned soil. Compound 1 was detected by HR-MS and MS/MS in extracts of all 6 isolates, and PCR amplification of genomic DNA from all 6 strains revealed the presence of the associated *rhlM* gene (Table S3).

Many methyltransferases, such as those of the isoprenylcysteine carboxyl methyltransferase family that includes RhlM, utilize S-adenosyl methionine (SAM) as a methyl donor. In order to further validate the biosynthetic origin of the carboxymethyl group of RLMEs, we performed a stable isotope labeling experiment using L-[D_3_]-methionine. Feeding L-[D_3_]-methionine to WT cultures led to incorporation of three deuterium atoms into RLME, as confirmed by detection of a +3 major isotopologue for each of the [M+NH_4_]^+^ and [M+Na]^+^ adducts via HR-MS/MS analysis of crude ethyl acetate extracts (Fig. S6). Notably, *P. caledonica* F3 also produces small quantities of O-desmethyl RLME (compound 5), and this mass feature did not show incorporation of any stable isotopes. Taken together, these results not only confirm that *rhlM* is responsible for rhamnolipid methylation, but further verify the activity of a SAM-dependent methyltransferase in RLME biosynthesis.

### *rhl* mutants display reduced antifungal activity

Having demonstrated the role of the genes *rhlA, rhlB,* and *rhlM* in RLME biosynthesis, we next sought to characterize the antifungal phenotype of each knockout mutant. Rhamnolipids (RLs) and HAAs, the respective major products that accumulate in Δ*rhlM* and Δ*rhlB* strains, are also known to exhibit antifungal activity (12–14). In order to compare the antifungal activity produced by these mutant strains against strains producing RLMEs, we grew the WT and the mutants in parallel under consistent growth conditions and performed a plug assay.

As expected, the mutant Δ*rhlA* displayed no antifungal activity, while knocking out the intermediate genes *rhlM* and *rhlB* reduced antifungal activity but did not completely abolish it (Fig 4A). Plugs taken from the Δ*rhlM* strain displayed a 26.0% reduction in average zone diameter, and plugs from Δ*rhlB* displayed a 95.7% reduction in average zone diameter. Complementation of the *rhlM*, *rhlB* or *rhlA* gene in the corresponding deletion mutant restored antifungal activity to WT levels, demonstrating that the observed phenotype is due to RLME production (Fig 4B). These data indicate that the inhibitory activity of *P. caledonica* F3 originates solely from products of the *rhl* pathway. Notably, comparison between WT and Δ*rhlM* suggests that carboxyl methylation significantly enhances antifungal activity of rhamnolipids.

**Figure 4.**
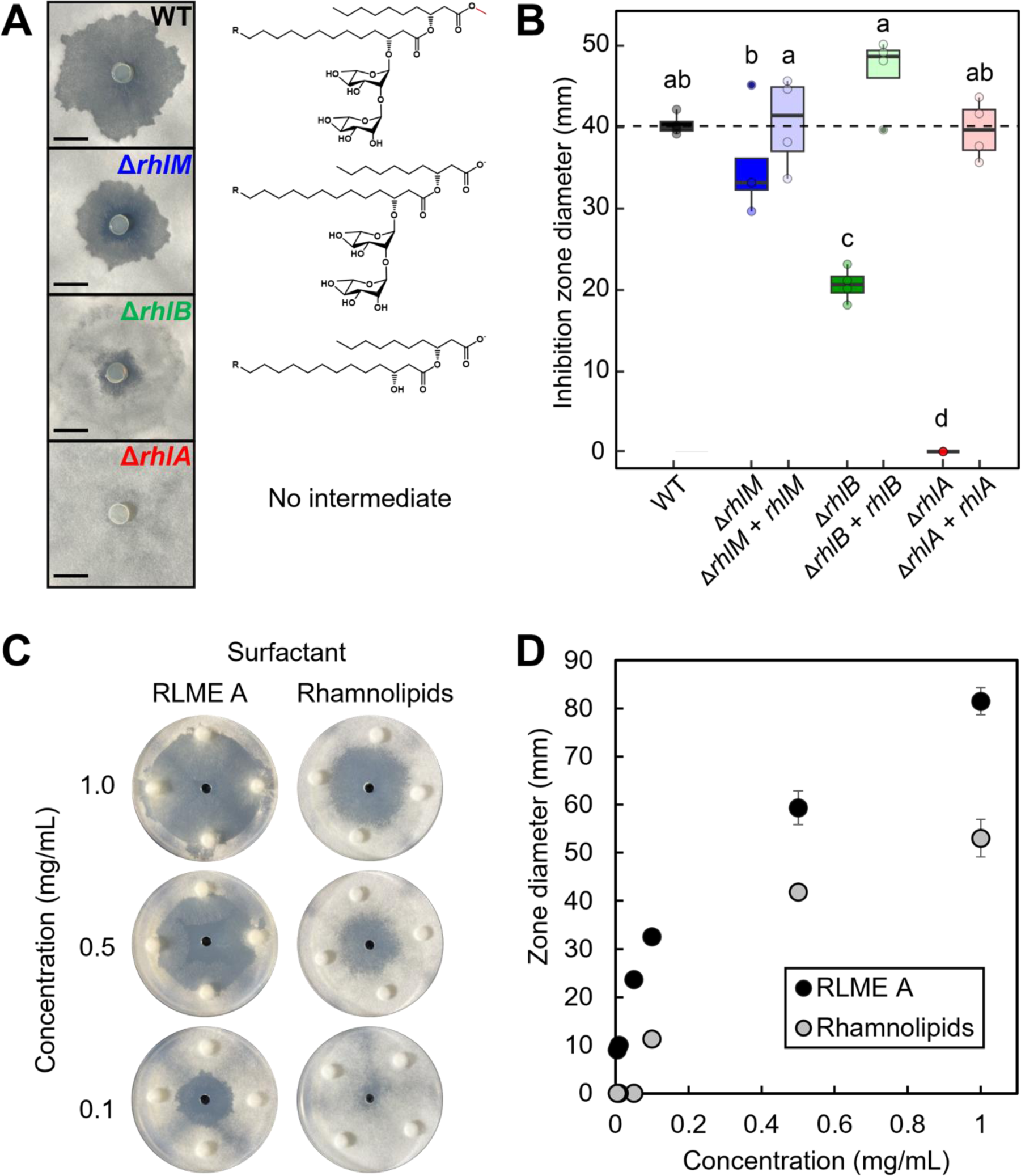
(A) *P. caledonica* F3 WT- or *rhl* mutant-conditioned plugs tested against *Pyronema omphalodes*. Images are representative of four biological replicates per strain and at least three independent experimental replicates. Structures to the right represent the major *rhl* pathway product for each strain. The RLME methyl group is highlighted in red. Scale bar is 1 cm. (B) Clearance zone diameters for *P. caledonica F3* WT, *rhl* mutant strains, and genetic complement strains tested against *Pyronema omphalodes*. Data are representative of four replicates per strain. Different letters indicate a statistically significant difference as determined by a one-way ANOVA and *post-hoc* Tukey’s test (*p* < 0.05). Dashed line indicates mean wildtype measurement. Inhibition of *Pyronema omphalodes* using different concentrations of purified rhamnolipid methyl ester A (RLME A) from *Paraburkholderia sp. F3* or Rhamnolipids (Di-Rhamnolipid dominant mixture) from *Pseudomonas aeruginosa* (Sigma-Aldrich). Compounds were solubilized and diluted in methanol and 15 μL of each solution was applied in the central well. Images are representative of three technical replicates. (D) Quantification of the inhibition zone diameters from tested RLME A and Rhamnolipids mixture. Points represent the mean of three replicates for each condition, and error bars represent standard deviation.

### Rhamnolipid methyl ester A is a stronger antifungal than rhamnolipids from Pseudomonas aeruginosa

Although the agar plug assays above suggest that RLMEs are stronger antifungals than unmethylated rhamnolipids, live cell cultures may exhibit variability in both metabolite concentrations and compositions. This variability complicates the analysis of structural differences and associated antifungal activity in WT and mutant strains. In order to circumvent this challenge, we assayed purified RLME A and a commercial standard of rhamnolipids (Di-Rhamnolipid dominant mixture) from *Pseudomonas aeruginosa* for antifungal activity against *P. omphalodes*, at concentrations ranging from 0.005 mg/mL to 0.1 mg/mL.

In all cases, RLME A was active at lower concentrations than the rhamnolipid mixture to create a zone of inhibition against *P. omphalodes*. Zones of inhibition are observed for RLME A at concentrations as low as 0.005 mg/mL, while a 20-fold higher concentration of 0.1 mg/mL was required for rhamnolipids to produce a zone of inhibition (Fig 4C). At equal concentrations, the zones of inhibition observed for RLME A were 41.8-186.8% larger than those observed for rhamnolipids (Fig. 4D). These findings indicate that RLME A has more potent antifungal activity than rhamnolipids.

### RLME and intermediates promote *P. caledonica* F3 motility *via* formation of a surfactant front

Several studies have linked rhamnolipid and HAA production with bacterial swarming motility in *Pseudomonas aeruginosa* and *Burkholderia* species (15–17). Specifically, evidence suggests that these molecules act as surfactants to promote swarm expansion (18). We sought to explore whether RLMEs and their precursors were connected to swarming motility of *P. caledonica* F3. We tested the impact of RLMEs on *P. caledonica* F3 motility in the WT and in knockout mutants lacking *rhlM*, *rhlB*, and *rhlA*. We tested the motility phenotypes of these strains on a rich ISP2 medium containing 0.25% agar and on a minimal medium (MM) containing 0.5% agar. After incubation, we visualized the surfactant zones on MM + 0.5% agar plates using atomized oil spray (19).

We observed that the WT strain was motile and spread out radially on the agar surface with a pronounced undulate margin (Fig. 5A). Both Δ*rhlM* and Δ*rhlB* strains exhibited reduced motility, with 51.7% and 30.6% reductions in average swarming diameter, respectively. Furthermore, the Δ*rhlM* strain swarm edges were entire rather than undulate, while the Δ*rhlB* strain swarm edges were only slightly undulate. The atomized oil assay revealed that, like WT, Δ*rhlM* and Δ*rhlB* strains produced a surfactant front, but they were reduced in diameter by 9.4% and 9.0%, respectively (Fig. 5A). Deletion of *rhlA* completely abolished swarming motility, and no surfactant front was observed when Δ*rhlA* MM plates were sprayed with atomized oil. Reductions in swarming for each mutant were ameliorated with genetic complementation (Fig. 5C). These findings indicate that RLMEs and their precursors are directly involved in *P. caledonica* F3 swarming motility on agar surfaces *via* the formation of a surfactant front.

**Figure 5.**
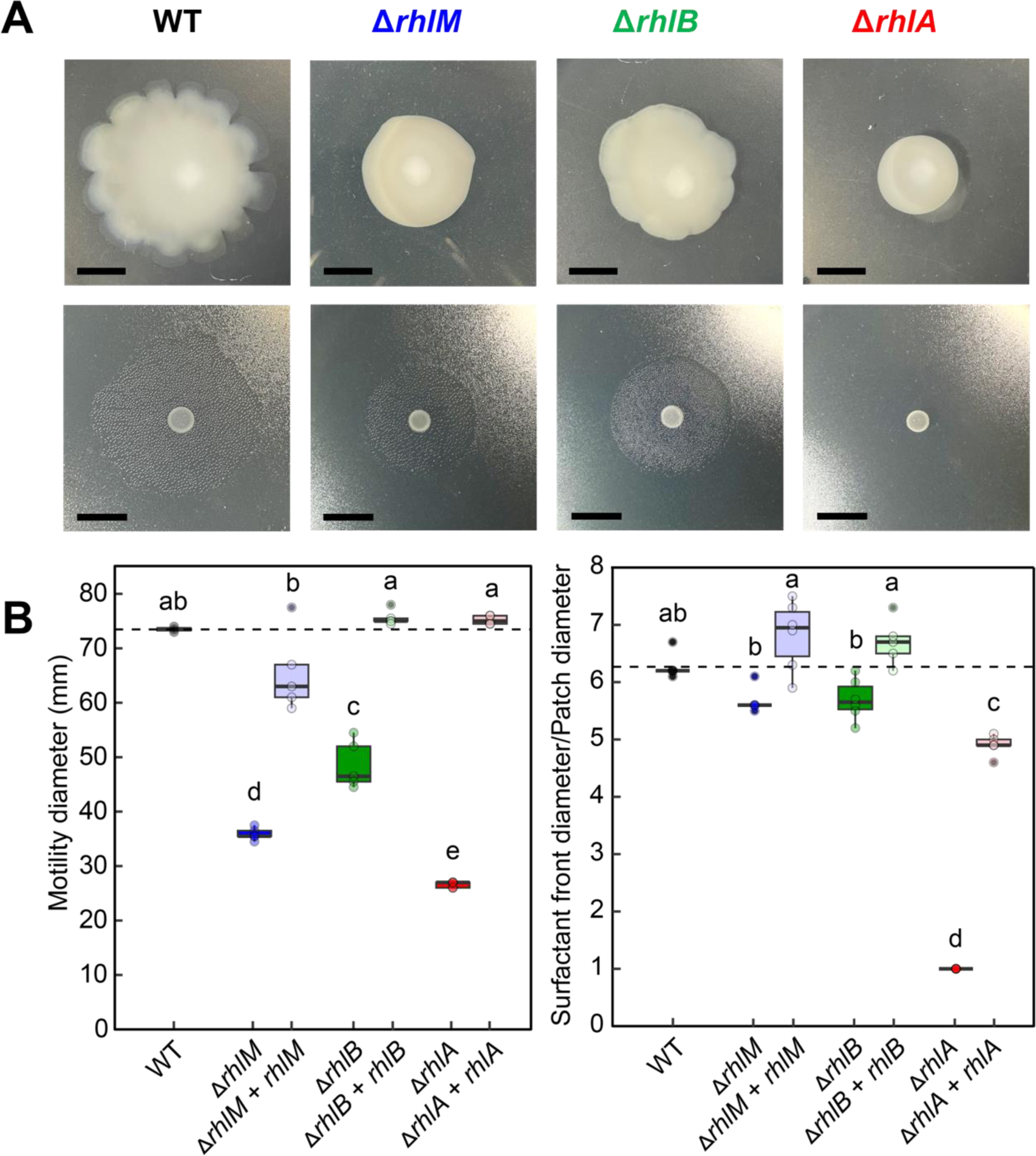
(A) Motility phenotypes (top) and surfactant zone production (bottom) for *P. caledonica F3* WT and *rhl* mutants. Images are representative of at least five biological replicates per strain and at least three independent experimental replicates. Scale bar is 1 cm. (B) Left: Swarming diameters for WT, *rhl* mutants, and genetic complement strains (WT swarming diameter normalized to 1). Right: Ratio of surfactant zone diameters to patch diameter for WT, *rhl* mutants, and genetic complement strains. Different letters indicate a statistically significant difference as determined by a one-way ANOVA and *post-hoc* Tukey’s test (*p* < 0.05). Dashed line indicates mean wildtype measurement.

### Rhamnolipid methyl esters improve aqueous solubilization of PAH compounds

*P. caledonica* F3 was isolated from a burned soil environment, in which a substantial amount of carbon is present in the form of pyrolyzed organic matter (PyOM, or char). PyOM consists of a complex mixture of polycyclic aromatic hydrocarbons (PAHs), which are hydrophobic substrates and thus poorly bioaccessible (20). However, analysis of the *P. caledonica* F3 genome showed that it possesses full or partial pathways for degradation of benzoate, catechols, and other common intermediates of aerobic catabolism of aromatic substrates (Fig. S7-8). Previous studies have shown rhamnolipid-enhanced solubilization of hydrocarbons, such as alkanes (21, 22) and PAHs (23–25). Synthetic rhamnolipid methyl esters were also shown to further enhance hydrocarbon solubilization (22). Thus, we sought to investigate the ability of rhamnolipid methyl ester A (RLME A) to solubilize PAHs, and assess its performance alongside the commercial rhamnolipids standard.

We tested purified RLME A for solubilization of three PAHs (naphthalene, phenanthrene, and benzo[*a*]pyrene) in comparison to the rhamnolipids. Biosurfactant was supplied to the mixtures at a concentration exceeding literature values of critical micelle concentrations (CMC) for rhamnolipids, taking into account lower predicted CMC values for RLMEs (22). RLME A improved aqueous solubilization of each individual PAH by nearly 5-fold compared to base solubilities in water (Fig. 6), while rhamnolipids improved solubilization by only ∼2-fold. These data indicate that RLMEs significantly enhance the solubility of these PAHs to a greater extent than typical rhamnolipids.

**Figure 6.**
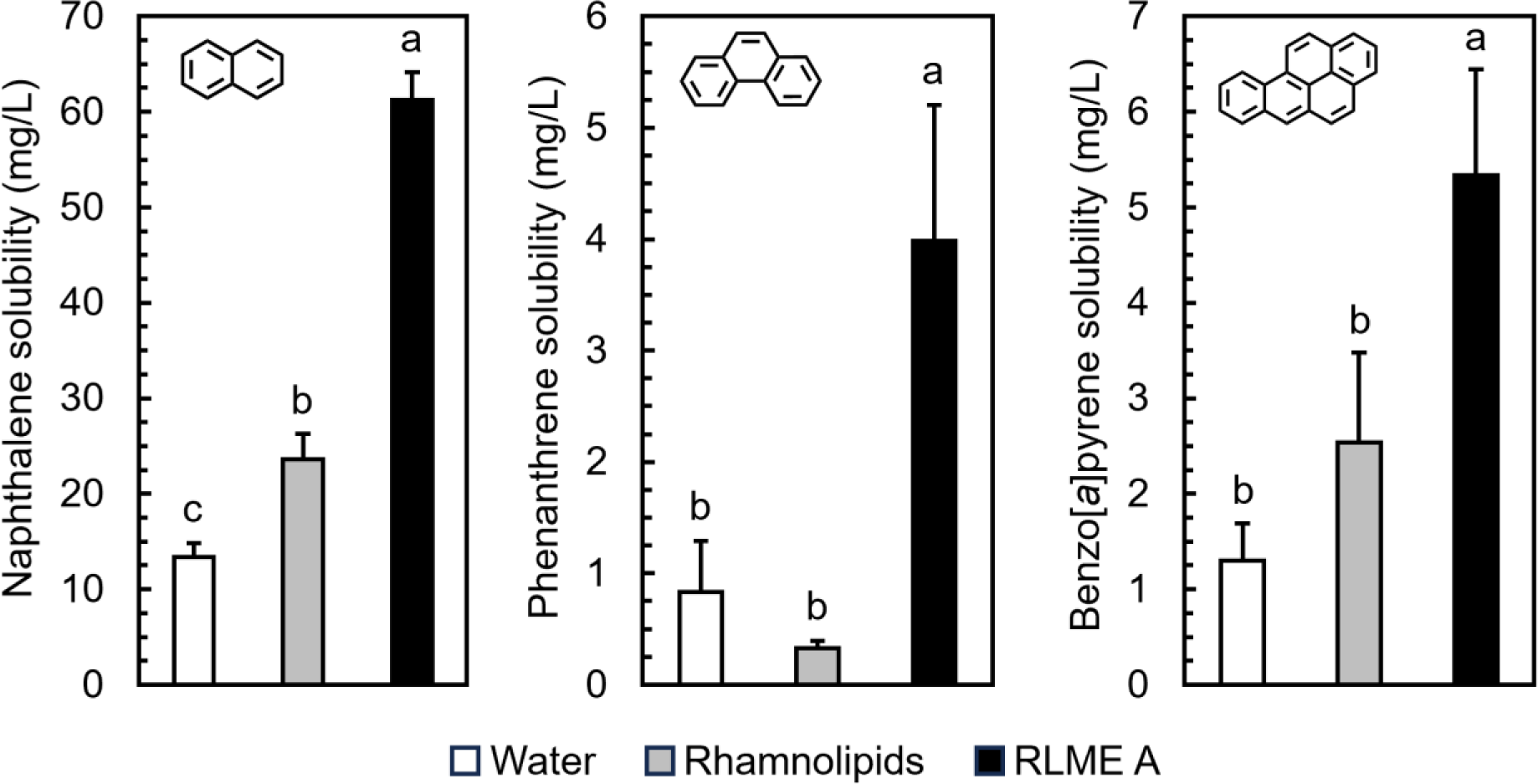
Solubilization of naphthalene (left), phenanthrene (center), and benzo[*a*]pyrene (right) in water supplemented with either 500 mg/L rhamnolipid methyl ester A (RLME A, (1)) from *P. caledonica* F3 or 500 mg/L Rhamnolipids (Di-Rhamnolipid dominant mixture, Sigma-Aldrich) from *Pseudomonas aeruginosa.* Different letters indicate a statistically significant difference as determined by a one-way ANOVA and *post-hoc* Tukey’s test (*p* < 0.05).

### RhlM is a putative SAM-utilizing integral membrane carboxyl methyltransferase

Multiple lines of investigation in this study point towards the carboxymethyl group in RLMEs as a key functional contributor to their surface activities, with relevance for antifungal activity, motility, and solubilization of potential PAH nutrient substrates. The *P. caledonica* F3 RhlM enzyme belongs to the isoprenylcysteine carboxyl methyltransferase (ICMT) family of integral membrane methyltransferases, which are predicted to accommodate both an amphiphilic substrate and the polar cofactor SAM. Aside from ICMTs, which are known to methylate the carboxyl group of prenylated protein substrates in eukaryotes, no other methyl acceptor substrates have been identified for proteins of this family. As a result, understanding the function of RhlM is of high interest. Thus, we sought to further explore RhlM with respect to its structure, substrate binding, and evolutionary context *in silico*.

In order to structurally characterize RhlM *in silico,* we generated a protein model for RhlM using AlphaFold (Fig. 7A, Fig. S10), which we aligned with the two crystallized members of the ICMT protein family, the archaeal Ma MTase (PDB: 4a2n) and eukaryotic *Tribolium castaneum* ICMT (PDB: 5vg9) (26, 27). The model of RhlM aligned well with both of these reported crystal structures, with pruned RMSD values of 1.118 Å and 0.793 Å, respectively (Fig. 7A, Table S5). After docking S-adenosyl homocysteine (SAH) into our RhlM model, we observed that SAH bound in similar conformations in RhlM and Ma MTase, as did SAM. Notably, two residues (E162 and H122) form interactions with SAM, closely resembling ligand interactions observed in SAH-bound Ma MTase and SAH-bound Tc ICMT (Fig. 7C, Fig. S11) (26, 27). Clustal Omega sequence alignments of RhlM with different ICMTs and other orthologs also demonstrated that these are conserved residues in the putative SAM cofactor binding pocket (Fig. S11).

**Figure 7.**
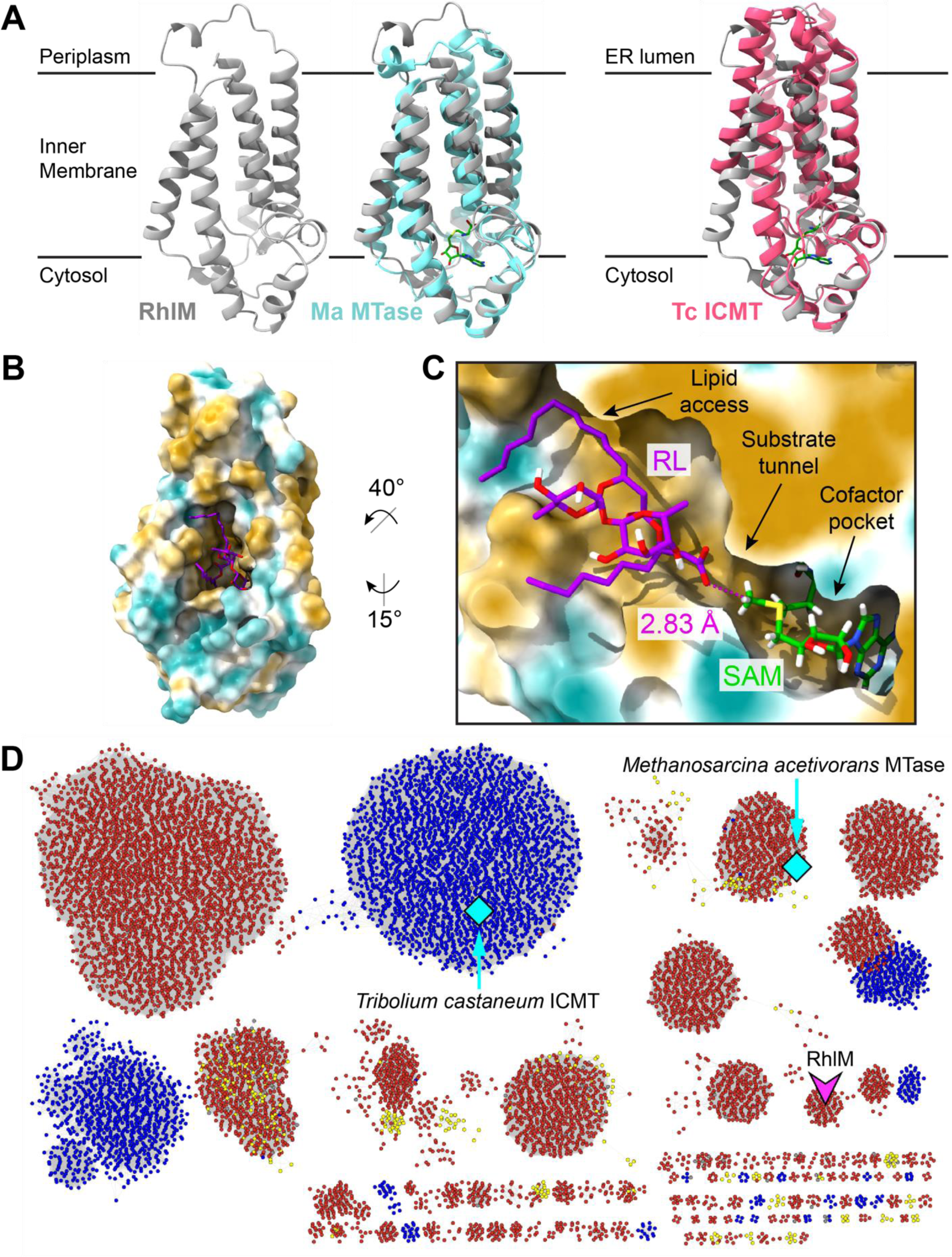
(A) AlphaFold model for RhlM (gray) alone, aligned with *Methanosarcina acetivorans* methyltransferase (PDB: 4a2n, cyan), and aligned with *Tribolium castaneum* ICMT (PDB: 5vg9, pink). SAH is shown in green. The membrane is indicated by two solid horizontal lines. (B) Surface representation of the rank 1 pose from Maestro ligand docking in RhlM, using an applied positional constraint for substrate carboxylate O and SAM methyl C. Docked pose was verified by molecular dynamics using Desmond. Surface is colored by molecular lipophilicity potential (gold = lipophilic, cyan = hydrophilic). (C) Cross-section of RhlM with substrate and cofactor bound. The upper hydrophobic region provides access for the lipid region of the RL substrate, while the lower region accommodates the carboxylate terminus, optimally positioning the carboxylate for methyl transfer from the SAM cofactor. Measured C-O distance is shown in magenta. (D) Sequence similarity network for ICMT protein family. Nodes are colored by domain: red, Bacteria; blue, Eukarya; yellow, Archaea; gray, unclassified. Magenta V-shape indicates *P. caledonica F3* RhlM, cyan diamonds indicate that a crystal structure is available. Clusters containing 3 or fewer nodes are omitted for clarity.

The experiments presented in Fig. 3A indicate that the major methyl accepting substrate of RhlM is likely the di-rhamnolipid Rha-Rha-C_14_-C_10_ (RL, compound 5) illustrated in Fig. 3C. To investigate binding of RL within RhlM, we performed sequential ligand docking experiments in Glide first with SAM followed by RL. We next performed a molecular dynamics simulation in Desmond, using RhlM modeled within a lipid membrane environment, in order to refine docked poses of RL within RhlM. These simulations place the RL lipid chains in contact with hydrophobic residues that form a lipid-binding region (Fig. 7C, Fig. S12). This lipid-binding pocket appears to be large and flexible enough to accommodate substrates with variable lipid chain lengths, in line with the observation of different RLME analogs produced by *P. caledonica* F3. This arrangement of compound 5 positions the RL carboxylate O within the active site in proximity to the SAM methyl C (∼3 Å) for methyl transfer (28). Taken together, these *in silico* results suggest a plausible mechanism for methylation of di-rhamnolipids by RhlM and set the stage for further structural and biochemical characterization of this novel enzyme.

In order to better define RhlM within the context of the ICMT protein family, we used the Enzyme Function Initiative’s Enzyme Similarity Tool (EFI-EST) (29) to generate a sequence similarity network (SSN) using all 14,240 sequences with the ICMT Pfam ID PF04140, with the RhlM sequence manually included. This network analysis revealed that the *P. caledonica* F3 RhlM sequence was part of a distinct cluster (Cluster 14) composed of RhlM homologs from predominantly *Burkholderia*, *Paraburkholderia*, and *Caballeronia* (Fig. 7C). Broadly, the full SSN comprises 230 clusters and is dominated by clusters of bacterial origin, with a few eukaryotic bacterial/archaeal subnetworks also present. This SSN analysis illustrates that ICMT paralogs are widespread, and found broadly in bacteria. These findings hint at a wide potential for future study within this enzyme class, with RhlM providing a starting point for elucidating the function of these enzymes across the bacterial domain.

## Discussion

The postfire soil environment is a challenging place for microbes to live and thrive - and one in which competition for limited resources is likely fierce as the microbial community successively assembles. Aside from biotic competition, postfire microbes face numerous other challenges, including increased hydrophobicity and poor nutrient solubility (30–33). However, our understanding of the mechanisms by which postfire microbes adapt to their perturbed environments remains limited.

Here, we report a set of rhamnolipid biosurfactants with an uncommon carboxymethylation, the rhamnolipid methyl esters (RLMEs), produced by a postfire bacterium, *Paraburkholderia caledonica* F3. These RLMEs inhibited the growth of a common pyrophilous fungus, promoted bacterial motility, and enhanced polycyclic aromatic hydrocarbon (PAH) solubility. In addition, we identified the genes required for RLME biosynthesis and found that a novel methyltransferase, termed RhlM, is required for the installation of the unusual carboxymethyl group found in these molecules. Notably, RhlM is the first characterized bacterial representative of a class of integral membrane methyltransferases that is widespread in bacteria. In sum, this study provides an example of specialized metabolism that may confer multiple adaptive advantages to postfire bacteria. Furthermore, this work highlights the potential in exploring perturbed environments for novel natural products and their associated enzymology.

### Probing perturbed environments leads to antifungal discovery

We employed an ecologically informed screening strategy intended to bias our findings towards unusual compounds. This approach began with isolates sourced from a postfire environment, which we cultivated using a medium containing an environmentally relevant carbon source in the form of pyrolyzed organic matter (PyOM). We screened for antifungal activity using an indicator strain, *Pyronema omphalodes*, which is found in high abundance in postfire soils (7, 34). Notably, we specifically looked for antifungal activity that was enhanced when the producing isolates were grown on PyOM, as opposed to a rich medium. Thus, at each step, we designed this strategy to increase the probability of identifying molecules that might carry ecological significance. This bioprospecting strategy led to the identification of the RLMEs. This approach represents a model framework to guide future discovery of bioactive compounds from perturbed environments, including molecules like RLMEs that possess modified chemical properties and/or function under unusual environmental conditions.

### Biosynthesis of RLMEs involves an integral membrane methyltransferase

Rhamnolipid methyl esters are members of a known class of rhamnolipid biosurfactants, which are thought to fulfill several different roles for their producing organisms. Rhamnolipid 1, first discovered from *Pseudomonas aeruginosa* (35), is known to exhibit antimicrobial properties and promote bacterial swarming motility (12–17). Rhamnolipids have also been identified as potential mediators for uptake of hydrophobic substrates by *P. aeruginosa* and consortia (22, 36–38). As a result of carboxyl methylation, the typically anionic lipid region of rhamnolipids is neutralized in RLMEs, rendering this region even more lipophilic. This key difference likely imparts stronger surface-active properties upon RLMEs, which may play a role in the enhanced antifungal activity, swarming motility, and PAH solubilization we demonstrate here for RLMEs from *P. caledonica* F3.

The enhanced surface activity of rhamnolipid esters has been previously established through studies that utilized synthetic RLMEs (22). However, to our knowledge, naturally occurring rhamnolipid methyl esters have been reported only once (39), without further verification of the biological origin of the appended methyl group. The identification of a natural producer of RLMEs, as well as elucidation of RLME biosynthesis, is therefore of notable interest for advancing the field of microbial biosurfactants as well as for industrial applications. We note that while an abundance of reports has focused on altering production titers (40–42), and structural components of rhamnolipids (43–45), none have provided a biological means toward esterification of the carboxy terminus. This work lays the foundation for development of heterologous expression or RhlM-dependent biocatalytic approaches for optimized production of RLMEs with altered chemical properties.

The rhamnolipid methyltransferase identified here, RhlM, is a member of a larger group of proteins known as the isoprenylcysteine carboxyl methyltransferase (ICMT) family. This family includes only two crystallized representatives; a eukaryotic ICMT (27) and an archaeal ortholog from *Methanosarcina acetivorans* (26), whose methyl-accepting substrate has not been determined. Thus, while some eukaryotic and archeal ICMT family proteins have been investigated, the functions of these proteins in bacteria remain unexplored. The designation of RhlM as a member of the ICMT protein family led us to further investigate the prevalence of these enzymes in bacteria. Our sequence similarity analysis of over 14,000 proteins revealed a large number of ICMT paralogs that form many discrete clusters across the bacterial domain.

The RhlM sequence was grouped as a part of a well-resolved cluster (Cluster 14). Genome neighborhood analysis of Cluster 14 sequences revealed that these genes all lie within rhamnolipid biosynthetic operons in the genomes of Betaproteobacteria, including those in the genera *Paraburkholderia, Burkholderia,* and *Caballeronia,* suggesting that RLME production is common within these clades. These results are in line with a recent bioinformatic survey that identified *rhl* clusters containing RhlM-like sequences across Burkholderiaceae genomes (46).

ICMT family proteins are integral membrane methyltransferases (47). Our modeling studies with RhlM highlight features of these enzymes that are relevant to their function in methylating amphiphilic substrates. For example, the lipid moieties of rhamnolipid intermediates synthesized in the cytosol are likely to partition to the inner membrane, where they have access to the lipid-binding tunnel leading toward the RhlM active site. This arrangement may facilitate the methylation of a wide array of amphiphilic substrates by other bacterial ICMTs, reflected by the manifold clusters of ICMT family proteins present in our sequence similarity analysis. In this regard, RhlM serves as a pivotal inroad into understanding the likely diverse functions of ICMT family enzymes in bacteria.

### Ecological implications for biosurfactant production within postfire soil communities

The array of challenges presented by postfire environments likely drive multifunctional adaptations in pyrophilous microbes. The results presented here support a model in which RLMEs are calibrated to function in postfire environments as mediators of interference competition, enhancers of motility, and potentiators of nutrient solubility.

The pyrophilous fungus *Pyronema omphalodes* is a dynamic member of the postfire microbial community, sometimes achieving relative abundances reaching 60.34% after fire before subsequently declining (7). This decline may be due to a host of factors, including interference competition mediated by chemical antagonism. This hypothesis underlied our choice to use *P. omphalodes* as an indicator organism in our screens to identify antimicrobials made by other postfire organisms. We found that RLMEs offered enhanced antifungal potency against *P. omphalodes* compared to typical rhamnolipids (Fig. 4). Whether this enhanced activity results from superior diffusibility on hydrophobic surfaces or other mechanisms, these results are consistent with the notion that RLMEs may have a direct function in interference competition among postfire community members.

Beyond competition, microbes face increased hydrophobicity owing to PyOM in burned soils, which likely limits dispersal to, and colonization of, new microniches (32). Here, we demonstrated that motility of *P. caledonica* F3 on agar surfaces was dependent on RLME production. This finding is broadly in line with previous studies showing that rhamnolipids enable swarming motility of *Pseudomonas* and *Burkholderia* species on similar surfaces (17, 48, 49). We hypothesize that production of enhanced biosurfactants like RLMEs may have strong implications for bacterial motility and colonization within hydrophobic burned soils. However, to our knowledge, swarming motility mediated by rhamnolipids has not been investigated in terrestrial environments. Efforts aimed at quantifying the advantages conferred by bacterial motility in burned soils are therefore of keen interest going forward.

Postfire microbes must contend with the additional challenge of nutrient limitation in burned soils, as a major component of the carbon is present in the form of polycyclic aromatic hydrocarbons (PAHs) associated with PyOM. These aromatic compounds are not only difficult to degrade, but their hydrophobicity decreases their bioavailability. Previous studies have shown that rhamnolipids can enhance the solubilization and degradation of PAHs (37, 50, 51). We hypothesized that RLMEs would likely exhibit increased ability to solubilize PAHs compared to typical rhamnolipids due to their methyl ester functionality. Our results indicate that indeed, RLMEs were significantly better at solubilizing three PAHs of varying complexity. While we observed greater solubility enhancement using RLMEs compared to rhamnolipids, these differences may be attributable to both carboxyl methylation as well as the 4-carbon longer acyl chain present in RLMEs from *P. caledonica* F3. Dissecting the relative contributions of these chemical modifications will require further investigation. While the ability of *P. caledonica* F3 to directly grow on PAHs was not assessed here, we note that the *P. caledonica* F3 genome features several dioxygenase genes likely involved in the oxidative catabolism of aromatic molecules (Fig. S7-8). Collectively, these results are consistent with a robust role for RLMEs in the local solubilization of PAHs which may serve as growth substrates for *P. caledonica* F3.

The multifunctional nature of RLMEs revealed by this work aligns with a growing body of evidence showing that many molecules initially regarded as antimicrobials may in fact play diverse roles for their producers, and may have variable impacts across microbial communities (8, 10, 52). A key recent example includes phenazines produced by *Pseudomonas* species that influence iron and phosphate bioavailability, but also exhibit antimicrobial activity(9). The possibility that RLMEs may serve as public goods for some community members by enhancing motility or nutrient access, while they may be detrimental others (*e.g. Pyronema, via* antifungal activity), warrants further exploration in the context of postfire community assembly and succession.

## Materials and Methods

### Strains and Growth Conditions

Bacterial strains and plasmids used are listed in Tables S1&2. Bacterial strains were cultured in ISP2 broth (malt extract 10 g/L, yeast extract 4 g/L, dextrose 4 g/L) or on ISP2 agar (ISP2 broth plus agar 18 g/L) unless otherwise indicated. Fungal strains were cultured on Vogel’s Minimal Medium agar (53). For atomized oil assays, *Paraburkholderia caledonica* F3 strains were spotted on a modified M9 minimal medium (20 mM ammonium chloride, 12 mM sodium phosphate dibasic, 22 mM potassium phosphate monobasic, 8.6 mM sodium chloride, 1 mM magnesium sulfate, 1 mM calcium chloride, 11 mM dextrose, 0.5% casamino acids) with 0.5% agar for atomized oil assays. *Paraburkholderia* sp. were initially isolated from burned soil samples collected from Blodgett Forest, using an enrichment method on PyOM agar (1.5 mM potassium phosphate monobasic, 4.7 mM ammonium chloride, 6.7 mM potassium chloride, 1 mM calcium chloride, 17 mM sodium chloride, 3 mM magnesium chloride, 20.6 mM sodium sulfate, 7.5 g/L noble agar, 0.5 g/L ground PyOM (Eastern White Pine wood pyrolyzed at 350 °C) (5), 7.5 µM iron(II) chloride, 0.8 µM cobalt(II) chloride, 0.5 µM manganese(II) chloride, 0.5 µM zinc(II) chloride, 0.1 µM boric acid anhydrous, 0.15 µM sodium molybdate dihydrate, 0.1 µM nickel(II) chloride, 11 nM copper(II) chloride, 58 nM 4-aminobenzoic acid, 8 nM D(+)-biotin, 164 nM nicotinamide, 42 nM D(+)-pantothenic acid hemicalcium, 83 nM pyridoxamine dihydrochloride, 59 nM thiamine hydrochloride, 37 nM cobalamin (Vitamin B12)). Initial screening of burned soil isolates was performed using ISP2 agar and PyOM agar.

### Antifungal screen for inhibition of Pyronema omphalodes

Bacterial isolates were cultured in liquid media for 2 days at 30 °C. Cells were pelleted, washed twice in PBS, resuspended in 500 μL 0.1% PBS, and spread on ISP2 agar or PyOM agar using sterile beads, in triplicate. Plates were incubated for five days at 30 °C. *Pyronema omphalodes* was grown on Vogel’s Minimal Medium (VMM) agar for three days at 25 °C. An agar plug from a plate of *P. omphalodes* was placed at the center of a fresh VMM agar plate, and agar plugs from each plate of bacteria were placed 2 cm from *P. omphalodes*. The presence (a zone of clearing) or absence (fungal growth around the agar plug) of antifungal activity was recorded after three days at 25 °C. Measurements of zones of clearance were recorded. Agar plugs from uninoculated agar plates were used as negative controls. Chemical extracts and fractions were resuspended in methanol to an approximate concentration of 1 mg/mL. Purified compounds and standards were resuspended in methanol and diluted to the appropriate concentration. For all solutions, 15 μL was spotted in a 1 cm-wide circular well.

### Chemical extractions

Ten 5 mm plugs were collected from culture plates and placed in a 2 mL microtube. Plugs were extracted with 750 µL ethyl acetate, sonicated for 10 min, and left at room temperature for 1 h. Tubes were centrifuged to pellet biomass, and extract was transferred to a new microtube and dried under vacuum at 45 °C. An extraction control of sterile ISP2 agar plates was performed in parallel.

### LC-HRMS and LC-MS/MS Analysis

Samples were analyzed by a Ultra-High Pressure Liquid Chromatography (UHPLC) system (Dionex Ultimate 3000, ThermoFisher, USA) coupled to a high resolution mass spectrometer (HRMS, Thermo Q-Exactive Quadrupole-Orbitrap, ThermoFisher, USA) using a Heated Electrospray ionization (HESI) source, using a C18 column (50 mm x 2.1 mm, 2.2 μm, Thermo Scientific AcclaimTM RSLC). Unless otherwise specified, the UHPLC method was as follows: 0-1 minute 10% ACN + 0.1% FA, a gradient from 1-11 minutes of 10% to 99% ACN + 0.1% FA, 11-14.5 minutes of 99% ACN + 0.1% FA and re-equilibration of the column back into 10% ACN + 0.1% FA from 14.5-18 minutes, injection volume of 5 μL, flow rate of 0.4 mL/min and column oven at 35°C. The full MS1 scan was performed in positive mode, resolution of 35,000 full width at half-maximum (FWHM), automatic gain control (AGC) target of 1 x 10e6 ions and a maximum ion injection time (IT) of 100 ms, mass range from m/z 200-2000. MS/MS analysis was acquired using a data-dependent Top5 method at a resolution of 17,500 FWHM, AGC target of 1 x 10e5 ions and maximum ion IT of 50 ms, using an isolation window of 3 m/z and normalized collision energy (NCE) of 20, 30 and 45. Cone spray voltage was 3.5kV.

### Extraction and bioactivity-guided isolation of rhamnolipid methyl esters from Paraburkholderia caledonica F3

A 4 L culture of *Paraburkholderia caledonica F3* was grown for 5 days at 30 °C on 150 x 50 mm plates containing 40 mL of ISP2 agar medium. Agar was chopped and extracted twice in 1:1 ethyl acetate, sonicated for 10 min, and left at room temperature for 12 to 18 h. Extract was filtered using a Whatman 114V filter. The filtrate was dried with anhydrous sodium sulfate and concentrated using a rotary evaporator. The crude extract was partially purified by solid phase extraction (SPE) using a C-18 Sep-Pak® Vac 35 cc, 10 g cartridge (Waters). Extract was loaded onto the column in 50% methanol (MeOH) and eluted in increasing concentrations of 50%, 60%, and 100% MeOH. The fractions were tested for activity against *P. omphalodes*, and dried. Activity assays revealed the active compound to be in the 100% MeOH fraction. The 100% MeOH fraction was injected onto a C-18 column (250 mm x 4.6 mm, 5.0 µm, 120 Å; Thermo Scientific Acclaim RSLC) and analyzed *via* LC-MS/MS (1.0 mL/min flow) to optimize peak separation prior to semi-preparative HPLC purification.

Semi-preparative HPLC was run on a UHPLC system (Dionex UltiMate 3000, Thermo Fisher) with a C-18 column (250 mm x 10 mm, 5.0 µm, 120 Å; Thermo Scientific Acclaim RSLC). The 100% MeOH SPE fraction was dried and resuspended in 90% MeOH. HPLC was run at 5.9 mL/min flow with water (solvent A) and acetonitrile (solvent B) using the following gradient: 0-1 minute 90% B, a gradient from 1 to 11 minutes of 90% to 99% B, 11-15 minutes 99% B, a gradient from 15 to 15.5 minutes of 99% to 90% B, and re-equilibration of the column at 90% from 15.5-19.5 minutes. Multiple injections of 300 µL were run and fractions were pooled. Fractions were collected every 10 seconds from 8 to 15.5 min and dried in a SpeedVac (SPD-1010-115, Thermo Fisher). Rhamnolipid methyl esters did not exhibit a strong UV absorbance profile, and bioactivity-guided fractionation was paired with LC-MS/MS detection. Fractions 13-18 (11 to 12.25 min) exhibited activity and contained ions [M+NH^4^]^+^ = 738.4995 and [M+Na]^+^ = 743.4558.

### NMR characterization of rhamnolipid methyl ester A

The 1D and 2D NMR spectra of rhamnolipid methyl ester, including ^1^H NMR, ^1^H–^1^H COSY, ^1^H– ^13^C HSQC, and ^1^H–^13^C HMBC spectra, were acquired, respectively, on a Bruker Avance 900 NMR spectrometer (900 MHz for ^1^H and 225 MHz for ^13^C) equipped with a cryoprobe. For the NMR test, the sample was dissolved in DMSO-*d_6_* (Cambridge Isotope Laboratories, Inc.). Data were collected and reported as follows: chemical shift, integration multiplicity (s, singlet; d, doublet; t, triplet; m, multiplet), and coupling constant. Chemical shifts were reported using the DMSO-*d_6_* resonance as the internal standard for ^1^H-NMR DMSO-*d_6_*: *δ* = 2.50 *p.p.m.* and ^13^C-NMR DMSO-*d6*: *δ* = 39.5 *p.p.m*.

Rhamnolipid methyl ester A was isolated as white amorphous solid. The length of the two lipids chains were revealed by MS/MS analyses. The presence of two rhamnose groups was separately supported by their COSY correlations. The stereochemistry of these two rhamnose moieties was indicated by their proton-proton coupling constants. The connection between these two sugar rings was assigned as C-2’-O-C-1’’, based on ^3^*J*-HMBC correlations from H-2’ (*δ*_H_ 3.60) to C-1’’ (*δ*_C_ 101.7) and from H-1’’ (*δ*_H_ 4.78) to C-2’ (*δ*_C_ 98.3). The di-rhamnose was linked to C-14, following confirmed by ^3^*J*-HMBC correlations from H-14 (*δ*_H_ 3.86) to C-1’ (*δ*_C_ 98.3) and from H-1’ (*δ*_H_ 4.67) to C-2’ (*δ*_C_ 73.5). The configurations of C-3 and C-14 were both assigned as R based on the biosynthetic pathway.

### Stable isotope labeling experiments with D_3_-L-Methionine

*Paraburkholderia caledonica* F3 was grown in a minimal medium, containing 12.8 g/L Na_2_HPO_4_, 3.0 g/L KH_2_PO_4_, 0.5 g/L NaCl, 1.0 g/L NH_4_Cl, 4 g/L glucose, 2 mM MgSO_4_ solution, 0.1 mM CaCl_2_ solution, 10 mL/L of RPMI 1640 B vitamins mixture (Sigma) and 150 mg/L of each of the 20 amino acids, supplemented with or without (control) 1 mg/mL D_3_-labeled methionine. 2 mL liquid cultures were adjusted to an OD 600 of 0.05 using *P. caledonica F3* overnight cultures in the control medium and incubated at 30 °C with shaking at 200 rpm for 36 hours. Cultures were extracted in 1:1 ethyl acetate. The ethyl acetate layer was dried down and resuspended in 300 μL methanol, which was transferred to an LC-MS vial with a glass insert. Samples were analyzed by UHPLC-MS/MS as described above.

### Paraburkholderia caledonica F3 genome sequencing, assembly, and annotation

Genomic DNA was extracted from *P. caledonica* F3 that was grown in ISP2 for 18 h, using a modified phenol/chloroform extraction. DNA quality was confirmed using gel electrophoresis, Bioanalyzer, a NanoDrop One UV-Vis spectrophotometer, and a Qubit fluorometer (Invitrogen Qubit DNA-HS assay kit, Q32851). Genomic DNA was run through the UCB QB3 PacBio Sequel II sequencing pipeline. Raw reads were partitioned using seqtk and the genome was assembled *de novo* with Flye into three contigs at 989x coverage. The draft genome was annotated using the RASTtk pipeline (54). Genome data was deposited to JGI IMG-MER (accession number xxxxx) and NCBI (accession number xxxxx).

### Construction of *rhl* knockout strains and genetic complementation

Chromosomal gene deletions in *P. caledonica* F3 were generated *via* double allelic exchange mediated by a pEXG2-based suicide vector (55). For each gene deletion strain, a separate suicide vector was constructed containing an insert comprised of a ∼500 bp upstream homology region and a ∼500 bp downstream homology region from the target gene. Primers consisted of a (19-21 nt) binding region and a 20-nt 5’ overhang complementary to the other fragment (backbone or insert). Primers used to amplify the upstream and downstream regions of the target genes are listed in Table S2. The insert and PCR-amplified backbone were assembled via Gibson assembly (56), and transformed into *E. coli* DH5α (New England Biolabs) for plasmid selection. Single colonies were picked from LB gentamicin selection plates, and plasmids recovered using a miniprep kit (Bioneer AccuPrep® Plasmid Mini Extraction Kit). Recovered plasmids were sequenced at the UC Berkeley DNA Sequencing Facility to confirm no mutations in the homology region. Plasmids were electroporated into *P. caledonica* F3 and merodiploids were selected on LB gentamicin plates. Single colonies were streaked onto NSLB+15%(w/v) sucrose plates to counterselect for double-crossover events. True recombinants (gentamicin sensitive and sucrose resistant) were resolved from merodiploids (gentamicin resistant and sucrose sensitive) based on the loss of two vector-associated markers, gentamicin resistance (*aaC1*) and sucrose sensitivity (*sacB*). Plasmids for genetic complementation were constructed with Gibson assembly of a PCR-amplified gene into the pBBR1-MCS5 vector backbone (57).

### Swarming motility assay and atomized oil assay

Seed cultures of each strain were grown in ISP2 liquid medium overnight at 30 °C. After 16-20h of growth, cultures were then subcultured 1:20 into fresh liquid medium and grown to an OD of ∼0.6. Cells were pelleted, washed twice with PBS, and resuspended to an OD of 0.5 in PBS. 5 µL of each cell suspension was spotted at the center of a plate (ISP2 with 0.25% agar for swarming motility assays, or modified M9 minimal medium (20 mM ammonium chloride, 12 mM sodium phosphate dibasic, 22 mM potassium phosphate monobasic, 8.6 mM sodium chloride, 1 mM magnesium sulfate, 1 mM calcium chloride, 11 mM dextrose, 0.5% casamino acids) with 0.5% agar for atomized oil assays). Assays were performed with five replicates per strain.

Plates were incubated for 3 days at room temperature, then imaged using an iPhone 12S camera. For atomized oil assays, plates were sprayed with a thin layer of light mineral oil (Fisher Scientific) and imaged using an iPhone 12S camera with an oblique light source. Motility diameters and surfactant zone diameters were measured by randomly choosing two orthogonal lines and obtaining the average. Data was analyzed for statistical significance using a one-way ANOVA and *post-hoc* Tukey’s test (*p* < 0.05).

### PAH solubilization

0.25 mg of rhamnolipid methyl ester A (RLME A) or Rhamnolipids (95%, Di-Rhamnolipid dominant, Sigma-Aldrich) were aliquoted into 2-mL scintillation vials. Excess (∼10 mg) of either naphthalene, phenanthrene, or benzo[*a*]pyrene was measured into vials containing RLME A, Rhamnolipids, or nothing (base solubilization controls). Each treatment was tested in triplicate, except for the water and rhamnolipids conditions for phenanthrene, which were tested in duplicate. 500 µL of sterile water with 0.01 M sodium azide (to inhibit microbial growth) was added to each vial, which were then sonicated for 10 min. The mixtures were pipetted into 1.5-mL microcentrifuge tubes and centrifuged for 20 min at 15,000 xg to pellet the solids. Supernatants were transferred to clean glass vials and dried by SpeedVac. Samples were resuspended in 2x dichloromethane (DCM), diluted up to 2-fold, and measured using a UV-Vis spectrophotometer (Thermo Scientific Genesys 10S) at 228 nm (naphthalene), 254 nm (phenanthrene), and 295 nm (benzo[*a*]pyrene). A standard curve was constructed from solutions of naphthalene (2.5 to 15 mg/L), phenanthrene (0.5 to 2.5 mg/L), or benzo[*a*]pyrene (0.55 to 2.75 mg/L) in DCM (Fig. S9). Beer’s law was used to determine concentrations from absorbance readings using the standard curve. Statistical significance was determined using a one-way ANOVA and *post-hoc* Tukey’s test (*p* < 0.05).

### RhlM Ligand Docking and Molecular Dynamics

A protein model for RhlM was generated using AlphaFold through the LatchBio interface. Protein structure alignments were performed in Maestro. RhlM was aligned with Ma MTase (PDB ID: 4a2n), and the S-adenosyl homocysteine (SAH) ligand was extracted from Ma MTase and docked into the RhlM AlphaFold model using Glide. This RhlM-SAH structure was then used to define the receptor grid for docking S-adenosyl methionine (SAM). The RhlM-SAM structure was used to dock Rha-Rha-C_14_-C_10_ (RL, compound 5) within a defined receptor grid and with an applied positional constraint of 4 Å between the RL carboxylate O and the SAM methyl C. Molecular dynamics simulations were performed using Desmond with a modeled lipid membrane environment (DPPC), pre-equilibrated at 325 K. Residues 4-24, 35-55, 76-94, and 144-163 were manually selected for inclusion in membrane-bound region. Multiple sequence alignments were performed using Clustal Omega. Structural alignments of RhlM with *Methanosarcina acetivorans* MTase (PDB: 4a2n) and *Tribolium castaneum* ICMT (PDB: 5vg9) were performed using ChimeraX (58).

### Sequence similarity and genome neighborhood analysis

A colored SSN was generated using the EFI-EST SSN tool for the ICMT family using the Pfam PF04140 with the sequence for RhlM added (29). An alignment score of 35 and sequence length cutoffs of no shorter than 150 and no longer than 330 amino acids was used. This SSN was visualized and analyzed using Cytoscape (59). The EFI-EST genome neighborhood tool (GNT) was then applied to this SSN using standard parameters to calculate the GND. Operons within the cluster containing RhlM (Cluster 14) were visually inspected in the GND explorer.

## Acknowledgments

We thank Kathy Durkin of the Molecular Graphics and Computational Facility (MGCF) at UC Berkeley for guidance with ligand docking and molecular dynamics experiments. The MGCF is supported by National Institutes of Health (NIH) award S10OD034382. We thank Steven Lindow for technical assistance with the atomized oil assay, and for helpful discussions. We are grateful to Eric Neubauer Vickers and Adina Lewis for assistance in purification of rhamnolipid methyl ester A. We also thank Neem Patel for isolation of *Paraburkholderia* strains from soil, Tom Bruns for isolation of *Pyronema omphalodes*, Thea Whitman for PyOM material, and Josephine Chandler and Max Sosa for plasmids pBBR1MCS-5 and pEXG2. This work was supported by Department of Energy (DOE) DE-SC0020351 and NIH R35GM128849 awarded to M.F.T. M.D.L. acknowledges the support of a National Science Foundation Graduate Research Fellowship. W.Z. and Y.D. were supported by NIH R01GM136758. Molecular graphics and analyses performed with UCSF ChimeraX, developed at the University of California, San Francisco, with support from NIH R01-GM129325 and the Office of Cyber Infrastructure and Computational Biology, National Institute of Allergy and Infectious Diseases.

## References

1. G. Certini, Effects of fire on properties of forest soils: a review. Oecologia 143, 1–10 (2005).

2. C. Santín, S. H. Doerr, C. M. Preston, G. González-Rodríguez, Pyrogenic organic matter production from wildfires: a missing sink in the global carbon cycle. Glob Chang Biol 21, 1621–1633 (2015).

3. J. E. Keeley, J. G. Pausas, P. W. Rundel, W. J. Bond, R. A. Bradstock, Fire as an evolutionary pressure shaping plant traits. Trends Plant Sci 16, 406–411 (2011).

4. N. C. Dove, N. Taş, S. C. Hart, Ecological and genomic responses of soil microbiomes to high-severity wildfire: linking community assembly to functional potential. ISME J 16, 1853–1863 (2022).

5. M. S. Fischer, et al., Pyrolyzed Substrates Induce Aromatic Compound Metabolism in the Post-fire Fungus, Pyronema domesticum. Frontiers in Microbiology 12 (2021).

6. D. Ghosal, S. Ghosh, T. K. Dutta, Y. Ahn, Current State of Knowledge in Microbial Degradation of Polycyclic Aromatic Hydrocarbons (PAHs): A Review. Frontiers in Microbiology 7 (2016).

7. T. D. Bruns, J. A. Chung, A. A. Carver, S. I. Glassman, A simple pyrocosm for studying soil microbial response to fire reveals a rapid, massive response by Pyronema species. PLoS One 15, e0222691 (2020).

8. D. L. McRose, D. K. Newman, Redox-active antibiotics enhance phosphorus bioavailability. Science (1979) 371, 1033–1037 (2021).

9. D. L. McRose, J. Li, D. K. Newman, The chemical ecology of coumarins and phenazines affects iron acquisition by pseudomonads. Proceedings of the National Academy of Sciences 120, e2217951120 (2023).

10. K. O. Thalhammer, D. K. Newman, A phenazine-inspired framework for identifying biological functions of microbial redox-active metabolites. Curr Opin Chem Biol 75, 102320 (2023).

11. J. Bauer, K. Brandenburg, U. Zähringer, J. Rademann, Chemical Synthesis of a Glycolipid Library by a Solid-Phase Strategy Allows Elucidation of the Structural Specificity of Immunostimulation by Rhamnolipids. Chemistry – A European Journal 12, 7116–7124 (2006).

12. M. E. Stanghellini, R. M. Miller, BIOSURFACTANTS: Their Identity and Potential Efficacy in the Biological Control of Zoosporic Plant Pathogens. Plant Dis 81, 4–12 (1997).

13. S. Lang, E. Katsiwela, F. Wagner, Antimicrobial Effects of Biosurfactants. Lipid / Fett 91, 363–366 (1989).

14. D. Li, W. Tao, D. Yu, S. Li, Emulsifying Properties of Rhamnolipids and Their In Vitro Antifungal Activity against Plant Pathogenic Fungi. Molecules 27 (2022).

15. E. Déziel, F. Lépine, S. Milot, R. Villemur, rhlA is required for the production of a novel biosurfactant promoting swarming motility in Pseudomonas aeruginosa: 3-(3-hydroxyalkanoyloxy)alkanoic acids (HAAs), the precursors of rhamnolipids. Microbiology (N Y) 149, 2005–2013 (2003).

16. D. Dubeau, E. Déziel, D. E. Woods, F. Lépine, Burkholderia thailandensis harbors two identical rhl gene clusters responsible for the biosynthesis of rhamnolipids. BMC Microbiol 9, 263 (2009).

17. N. C. Caiazza, R. M. Q. Shanks, G. A. O’Toole, Rhamnolipids Modulate Swarming Motility Patterns of Pseudomonas aeruginosa. J Bacteriol 187, 7351–7361 (2005).

18. D. B. Kearns, A field guide to bacterial swarming motility. Nat Rev Microbiol 8, 634–644 (2010).

19. A. Y. Burch, B. K. Shimada, P. J. Browne, S. E. Lindow, Novel High-Throughput Detection Method To Assess Bacterial Surfactant Production. Appl Environ Microbiol 76, 5363–5372 (2010).

20. A. R. Johnsen, U. Karlson, Diffuse PAH contamination of surface soils: environmental occurrence, bioavailability, and microbial degradation. Appl Microbiol Biotechnol 76, 533– 543 (2007).

21. Y. Zhang, R. M. Miller, Enhanced octadecane dispersion and biodegradation by a Pseudomonas rhamnolipid surfactant (biosurfactant). Appl Environ Microbiol 58, 3276– 3282 (1992).

22. Y. Zhang, R. M. Miller, Effect of Rhamnolipid (Biosurfactant) Structure on Solubilization and Biodegradation of n-Alkanes. Appl Environ Microbiol 61, 2247–2251 (1995).

23. H. Yu, G. Huang, J. Wei, C. An, Solubilization of Mixed Polycyclic Aromatic Hydrocarbons through a Rhamnolipid Biosurfactant. J Environ Qual 40, 477–483 (2011).

24. S. Li, et al., Effect of rhamnolipid biosurfactant on solubilization of polycyclic aromatic hydrocarbons. Mar Pollut Bull 101, 219–225 (2015).

25. C.-H. Kim, et al., Desorption and solubilization of anthracene by a rhamnolipid biosurfactant from Rhodococcus fascians. Water Environment Research 91, 739–747 (2019).

26. J. Yang, et al., Mechanism of Isoprenylcysteine Carboxyl Methylation from the Crystal Structure of the Integral Membrane Methyltransferase ICMT. Mol Cell 44, 997–1004 (2011).

27. M. M. Diver, L. Pedi, A. Koide, S. Koide, S. B. Long, Atomic structure of the eukaryotic intramembrane RAS methyltransferase ICMT. Nature 553, 526–529 (2018).

28. D. Tsao, L. Diatchenko, N. V Dokholyan, Structural Mechanism of S-Adenosyl Methionine Binding to Catechol O-Methyltransferase. PLoS One 6, e24287-(2011).

29. R. Zallot, N. Oberg, J. A. Gerlt, The EFI Web Resource for Genomic Enzymology Tools: Leveraging Protein, Genome, and Metagenome Databases to Discover Novel Enzymes and Metabolic Pathways. Biochemistry 58, 4169–4182 (2019).

30. S. H. Doerr, A. D. Thomas, The role of soil moisture in controlling water repellency: new evidence from forest soils in Portugal. J Hydrol (Amst) 231–232, 134–147 (2000).

31. J. A. Moody, D. A. Martin, S. L. Haire, D. A. Kinner, Linking runoff response to burn severity after a wildfire. Hydrol Process 22, 2063–2074 (2008).

32. M. Gray, M. G. Johnson, M. I. Dragila, M. Kleber, Water uptake in biochars: The roles of porosity and hydrophobicity. Biomass Bioenergy 61, 196–205 (2014).

33. C. Gibson, et al., Weathering of pyrogenic organic matter induces fungal oxidative enzyme response in single culture inoculation experiments. Org Geochem 92, 32–41 (2016).

34. F. J. Seaver, Studies in Pyrophilous Fungi—I. The Occurrence and Cultivation of Pyronema. Mycologia 1, 131–139 (1909).

35. F. G. Jarvis, M. J. Johnson, A Glyco-lipide Produced by Pseudomonas Aeruginosa. J Am Chem Soc 71, 4124–4126 (1949).

36. S. S. Cameotra, P. Singh, Synthesis of rhamnolipid biosurfactant and mode of hexadecane uptake by Pseudomonas species. Microb Cell Fact 8, 16 (2009).

37. Q. Ruolin, X. Tao, J. Xiaoqiang, Engineering Pseudomonas putida To Produce Rhamnolipid Biosurfactants for Promoting Phenanthrene Biodegradation by a Two-Species Microbial Consortium. Microbiol Spectr 10, e00910–22 (2022).

38. R. A. Al-Tahhan, T. R. Sandrin, A. A. Bodour, R. M. Maier, Rhamnolipid-Induced Removal of Lipopolysaccharide from Pseudomonas aeruginosa: Effect on Cell Surface Properties and Interaction with Hydrophobic Substrates. Appl Environ Microbiol 66, 3262–3268 (2000).

39. T. Hirayama, I. Kato, Novel methyl rhamnolipids from Pseudomonas aeruginosa. FEBS Lett 139, 81–85 (1982).

40. T. Tiso, et al., Integration of Genetic and Process Engineering for Optimized Rhamnolipid Production Using Pseudomonas putida. Frontiers in Bioengineering and Biotechnology 8 (2020).

41. J. Beuker, et al., Integrated foam fractionation for heterologous rhamnolipid production with recombinant Pseudomonas putida in a bioreactor. AMB Express 6, 11 (2016).

42. T. Tiso, et al., Creating metabolic demand as an engineering strategy in Pseudomonas putida – Rhamnolipid synthesis as an example. Metab Eng Commun 3, 234–244 (2016).

43. N. Cabrera-Valladares, et al., Monorhamnolipids and 3-(3-hydroxyalkanoyloxy)alkanoic acids (HAAs) production using Escherichia coli as a heterologous host. Appl Microbiol Biotechnol 73, 187–194 (2006).

44. S. G. V. A. O. Costa, E. Déziel, F. Lépine, Characterization of rhamnolipid production by Burkholderia glumae. Lett Appl Microbiol 53, 620–627 (2011).

45. T. Tiso, et al., Designer rhamnolipids by reduction of congener diversity: production and characterization. Microb Cell Fact 16, 225 (2017).

46. M. Magri, A. M. Abdel-Mawgoud, Identification of putative producers of rhamnolipids/glycolipids and their transporters using genome mining. Curr Res Biotechnol 4, 152–166 (2022).

47. W. S. Yang, et al., Isoprenyl carboxyl methyltransferase inhibitors: a brief review including recent patents. Amino Acids 49, 1469–1485 (2017).

48. J. Yan, H. Monaco, J. B. Xavier, The Ultimate Guide to Bacterial Swarming: An Experimental Model to Study the Evolution of Cooperative Behavior. Annu Rev Microbiol 73, 293–312 (2019).

49. A. Nickzad, F. Lépine, E. Déziel, Quorum Sensing Controls Swarming Motility of Burkholderia glumae through Regulation of Rhamnolipids. PLoS One 10, e0128509 (2015).

50. S. Kumari, K. Gautam, M. Seth, S. Anbumani, N. Manickam, Bioremediation of polycyclic aromatic hydrocarbons in crude oil by bacterial consortium in soil amended with Eisenia fetida and rhamnolipid. Environmental Science and Pollution Research 30, 82517–82531 (2023).

51. Á. Martínez-Toledo, M. del Carmen Cuevas-Díaz, O. Guzmán-López, J. López-Luna, C. Ilizaliturri-Hernández, Evaluation of in situ biosurfactant production by inoculum of P. putida and nutrient addition for the removal of polycyclic aromatic hydrocarbons from aged oil-polluted soil. Biodegradation 33, 135–155 (2022).

52. L. E. P. Dietrich, T. K. Teal, A. Price-Whelan, D. K. Newman, Redox-Active Antibiotics Control Gene Expression and Community Behavior in Divergent Bacteria. Science (1979) 321, 1203–1206 (2008).

53. V. H. J., A convenient growth medium for Neurospora (medium N). Microbial Genet. Bull. 13, 42–43 (1956).

54. T. Brettin, et al., RASTtk: A modular and extensible implementation of the RAST algorithm for building custom annotation pipelines and annotating batches of genomes. Sci Rep 5, 8365 (2015).

55. L. R. Hmelo, et al., Precision-engineering the Pseudomonas aeruginosa genome with two-step allelic exchange. Nat Protoc 10, 1820–1841 (2015).

56. D. G. Gibson, et al., Enzymatic assembly of DNA molecules up to several hundred kilobases. Nat Methods 6, 343–345 (2009).

57. M. Kovach, R. Phillips, P. Elzer, R. M. I. Roop, K. M. Peterson, pBBR1MCS: a broad-host-range cloning vector. Biotechniques 16, 800–802 (1994).

58. E. F. Pettersen, et al., UCSF ChimeraX: Structure visualization for researchers, educators, and developers. Protein Science 30, 70–82 (2021).

59. P. Shannon, et al., Cytoscape: a software environment for integrated models of biomolecular interaction networks. Genome Res 13, 2498–2504 (2003).

