## Supplementary Tables and Figures for "Surface-active antibiotic production is a multifunctional adaptation for postfire microbes"

### Contents

**Supplementary Tables** 1

Supplementary Table 1. Strains used in this study
Supplementary Table 2. Oligonucleotide sequences used for plasmid construction and sequencing
Supplementary Table 3. Prevalence of RLMEs among burned soil *Paraburkholderia* isolates
Supplementary Table 4. NMR data of Compound 720 (RLME A) in DMSO-$d_6$
Supplementary Table 5. Protein structural alignment comparisons

**Supplementary Figures** 6

Supplementary Figure 1. $^1$H NMR spectrum of rhamnolipid methyl ester A, DMSO-$d_6$ at 900 MHz
Supplementary Figure 2. $^1$H-$^1$H COSY spectrum of rhamnolipid methyl ester, DMSO-$d_6$ at 900 MHz
Supplementary Figure 3. $^1$H-$^{13}$C HSQC spectrum of rhamnolipid methyl ester, DMSO-$d_6$ at 900 MHz
Supplementary Figure 4. $^1$H-$^{13}$C HMBC spectrum of rhamnolipid methyl ester, DMSO-$d_6$ at 900 MHz
Supplementary Figure 5. Extracted ion chromatograms for *P. caledonica* F3 strains
Supplementary Figure 6. Stable isotope labeling using $D_3$-Methionine
Supplementary Figure 7. Predicted pathways for toluene degradation in *P. caledonica* F3
Supplementary Figure 8. Predicted pathways for benzoate degradation in *P. caledonica* F3
Supplementary Figure 9. Standard curves for PAH solubilization experiments
Supplementary Figure 10. AlphaFold structure of RhlM colored by pLDDT score
Supplementary Figure 11. RhlM interactions with S-adenosyl methionine
Supplementary Figure 12. RhlM interactions with RL (compound 5)

***Supplementary Tables***

**Table S1.** Strains used in this study.

| Strain ID | Species/Strain Name | Relevant Genotype | Source |
|---|---|---|---|
| F3 | *Paraburkholderia caledonica* F3 | Wildtype | This study |
| ML59 | *Paraburkholderia caledonica* F3 | Δ*rhlM* | This study |
| ML65 | *Paraburkholderia caledonica* F3 | Δ*rhlB* | This study |
| ML91 | *Paraburkholderia caledonica* F3 | Δ*rhlA* | This study |
| ML77 | *Paraburkholderia caledonica* F3 | Δ*rhlM* + pBBR-MCS5-rhlM | This study |
| ML84 | *Paraburkholderia caledonica* F3 | Δ*rhlB* + pBBR-MCS5-rhlB | This study |
| ML137 | *Paraburkholderia caledonica* F3 | Δ*rhlA* + pBBR-MCS5-rhlA | This study |
| C1 | *Paraburkholderia kirstenboschensis* C2 | Wildtype | This study |
| C2 | *Paraburkholderia strydomiana* C1 | Wildtype | This study |
| A3-S | *Paraburkholderia kirstenboschensis* A3-S | Wildtype | This study |
| D1 | *Paraburkholderia kirstenboschensis* D1 | Wildtype | This study |
| D6-S | *Paraburkholderia kirstenboschensis* D6-S | Wildtype | This study |
| G6 | *Paraburkholderia kirstenboschensis* G6 | Wildtype | This study |
| G7-S | *Paraburkholderia kirstenboschensis* G7-S | Wildtype | This study |
| P1672 | *Pyronema omphalodes* | Wildtype | Tom Bruns |

Wild soil isolates were identified by closest 16S rRNA match on NCBI at the time of deposition to the collection. All *P. caledonica* F3 knockout and complementation strains were generated using primers in Table S2.

**Table S2.** Oligonucleotide sequences used for plasmid construction and sequencing.

| Name | Sequence (5' --> 3') |
|---|---|
| 16S-27F | AGAGTTTGATCCTGGCTCAG |
| 16S-1492-R | GGTTACCTTGTTACGSCTT |
| **Primers for gene deletion** | |
| RhlM-Up-F | aagcttctgcaggtcgactcCGATGCTCGCCTATGTGACCC |
| RhlM-Up-R | gacctcaatagagCGTGTGGTAGATCGTCATTTTCAG |
| RhlM-Down-F | atgacgatctaccacacgCTCTATTGAGGTCCGCGCG |
| RhlM-Down-R | gagcccggggatcctctaAGCACGCCTTTCTCGATCATTTC |
| pEXG2-RhlM-Gibson-F | aaatgatcgagaaaggcgtgctTAGAGGATCCCCGGGCTCG |
| pEXG2-RhlM-Gibson-R | ggtcacataggcgagcatcgGAGTCGACCTGCAGAAGCTTGC |
| RhlM-Seq-F | TCATTGCTCGCCCTGGCG |
| RhlM-Seq-R | GGCCGAGGATACGGCTTG |
| pEXG2-seq-F | tgttgcatgggcataaagttg |
| pEXG2-seq-R | tcaacgacaggagcacgatc |
| RhlB-Up-F | ttccacacattatacgagccggaagcataaatgtaaagcaAAGCGTCCTCCTGGGCGAAG |
| RhlB-Up-R | gttcaggcgacTGCGGTGATGACGATTTGTGC |
| RhlB-Down-F | aatcgtcatcaccgcaGTCGCCTGAACGAATCCGG |
| RhlB-Down-R | taaggtaccgaattcgagctcgagcccggggatcctctagCTGGTCGATGATCCAGCCGC |
| RhlB-Seq-F | GTCTGGCTCGTCTACTGG |
| RhlB-Seq-R | AAAGTGCTGCGCATGCGG |
| pEXG2-RhlB-Gibson-F | gcggctggatcatcgaccagCTAGAGGATCCCCGGGCTC |
| pEXG2-RhlB-Gibson-R | cttcgcccaggaggacgcttTGCTTTACATTTATGCTTCCGGC |
| RhlA-Up-F | caagcttctgcaggtcgactcGTCTTGCGGCCTAGGTGTC |
| RhlA-Up-R | ccgaaatcggtgttTTCGACGGACATAGAGCCCC |
| RhlA-Down-F | tctatgtccgtcgaaAACACCGATTTCGGAGGCTG |
| RhlA-Down-R | cgagcccggggatcctctaTACGGGTCGCCGAAATACTCG |
| RhlA-Seq-F | ACATACCCGCGACCGGC |
| RhlA-Seq-R | TCG CGC GGA CCT CAA TAG |
| pEXG2-RhlA-Gibson-F | cgagtatttcggcgacccgtaTAGAGGATCCCCGGGCTCG |
| pEXG2-RhlA-Gibson-R | gacacctaggccgcaagacGAGTCGACCTGCAGAAGCTTGC |
| **Primers for genetic complementation constructs** | |
| RhlM-F | aattcgatatcaagcttatcgGAGTCGCGCGGACCTCAATAG |
| RhlM-R | ccctcgaggtcgacggtatTTGAACACCGATTTCGGAGGCTG |
| BB-RhlM-F | gcctccgaaatcggtgttcaaATACCGTCGACCTCGAGGG |
| BB-RhlM-R | attgaggtccgcgcgactcCGATAAGCTTGATATCGAATTCCTGC |
| RhlB-F | aattcgatatcaagcttatcgTCAGGCGACCGAACGCGTG |
| RhlB-R | cccctcgaggtcgacggtatATTGAGGTCCGCGCGACTC |
| BB-RhlB-F | tgagtcgcgcggacctcaatATACCGTCGACCTCGAGGG |
| BB-RhlB-R | cacgcgttcggtcgcctgaCGATAAGCTTGATATCGAATTCCTGC |
| RhlA-F | cccctcgaggtcgacggtatCATCCACCGTAACATCCTGG |
| RhlA-R | gcaggaattcgatatcaagcttatcgCAAGCGTGTGGTAGATCGTC |
| BB-RhlA-F | gacgatctaccacacgcttgCGATAAGCTTGATATCGAATTCCTGC |
| BB-RhlA-R | ccaggatgttacggtggatgATACCGTCGACCTCGAGGGG |
| pBBR1MCS5-Seq-R | CAGGAAACAGCTATGACC |
| pBBR1MCS5-Seq-F | TGTAAAACGACGGCCAGT |

F = forward, R = reverse. Lowercase letters represent Gibson homology.

**Table S3.** Prevalence of rhamnolipid methyl esters among different *Paraburkholderia* burned soil isolates.

| Strain | Species | Source | | | *Pyronema* inhibition | HR-MS, MS/MS | *rhlM* |
|---|---|---|---|---|---|---|---|
| | | Plot | Depth | Collection date | | | |
| F3 | *P. caledonica* | 321E | 3-6 cm | 20-Nov-18 | + | + | + |
| A3-S | *P. kirstenboschensis* | 321E | 3-6 cm | 29-Oct-18 | + | + | + |
| G7-S | *P. kirstenboschensis* | 321E | 3-6 cm | 27-Nov-18 | + | + | + |
| C2 | *P. strydomiana* | 240 | 0-10 cm | 12-Oct-18 | + | + | + |
| G6 | *P. kirstenboschensis* | 321E | 0-10 cm | 16-Oct-18 | + | + | + |
| D1 | *P. ginsengisoli* | 240 | 0-10 cm | 12-Oct-18 | + | + | + |
| D6-S | *P. kirstenboschensis* | 321E | 0-3 cm | 29-Oct-18 | + | + | + |

The prevalence of RLMEs was examined using HR-MS and MS/MS fragmentation in addition to PCR. Positive data indicate that inhibition of *Pyronema omphalodes* P1672 was observed, RLMEs were detected in culture extracts, and *rhlM* was amplified from gDNA. Plot 321E was burned on October 16, 2018, and Plot 240 was an unburned plot. Depth and Collection date refer to the collected soil samples from which the strain was subsequently isolated. Species identification is based on a BLAST search of the 16S PCR amplicon and a >97% similarity cutoff.

**Table S4.** NMR Data of Compound 720 (Rhamnolipid methyl ester A) in DMSO-d6.

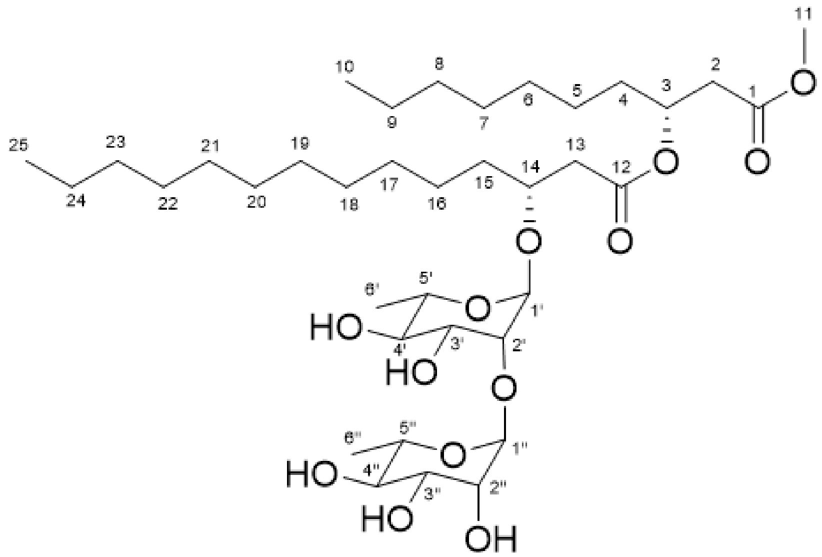

| Position | $\delta_H$ (*J* in Hz) | $\delta_C$ (C type) | Position | $\delta_H$ (*J* in Hz) | $\delta_C$ (C type) |
|---|---|---|---|---|---|
| 1 | | 170.4, C | 24 | 0.84 d 7.0 | 13.7, $CH_3$ |
| 2 | 2.63 dd 15.6, 5.5 | 38.1, $CH_2$ | 1’ | 4.67 brs | 98.3, CH |
| | 2.56 dd 15.6, 7.7 | | 2’ | 3.60 brs | 76.5, CH |
| 3 | 5.08 m | 69.7, CH | 3’ | 3.47 d 11.9 | 70.1, CH |
| 4 | 1.54 m | 33.0, $CH_2$ | 4’ | 3.17 dd 11.9, 11.9 | 71.7, CH |
| 5 | 1.23 m | 24.1, $CH_2$ | 5’ | 3.46 m | 68.5, CH |
| 6 | 1.23 m | 28.7, $CH_2$ | 6’ | 1.11 d 6.2 | 17.5, $CH_3$ |
| 7 | 1.23 m | 28.7, $CH_2$ | | | |
| 8 | 1.22 m | 31.1, $CH_2$ | | | |
| 9 | 1.26 m | 21.8, $CH_2$ | | | |
| 10 | 0.85 d 7.0 | 13.7, $CH_3$ | 1’’ | 4.78 brs | 101.7, CH |
| 11 | 3.58 s | 51.1, $CH_3$ | 2’’ | 3.69 brs | 69.9, CH |
| 12 | | 169.9, C | 3’’ | 3.39 dd 9.5, 3.3 | 70.3, CH |
| 13 | 2.48 dd 15.0, 5.9 | 39.8, $CH_2$ | 4’’ | 3.17 dd 9.5, 9.5 | 71.7, CH |
| | 2.44 dd 15.0, 6.6 | | 5’’ | 3.43 m | 68.4, CH |
| 14 | 3.86 m | 73.5, CH | 6’’ | 1.09 d 6.2 | 17.5, $CH_3$ |
| 15 | 1.44 m | 32.6, $CH_2$ | | | |
| 16 | 1.23 m | 24.4, $CH_2$ | | | |
| 17-22 | 1.23 m | 28.7, $CH_2$ | | | |
| 23 | 1.22 m | 31.1, $CH_2$ | | | |

**Table S5.** Protein structural alignment comparisons of RhlM AlphaFold model with ICMT family crystal structures.

| | PDB: 4a2n | PDB: 5vg9 |
|---|---|---|
| Root mean square deviation (RMSD), pruned | 1.118 Å | 0.793 Å |
| RMSD, all | 4.445 Å | 8.275 Å |
| Sequence alignment score | 304.1 | 255.7 |

Structural alignments performed in ChimeraX.

***Supplementary Figures***

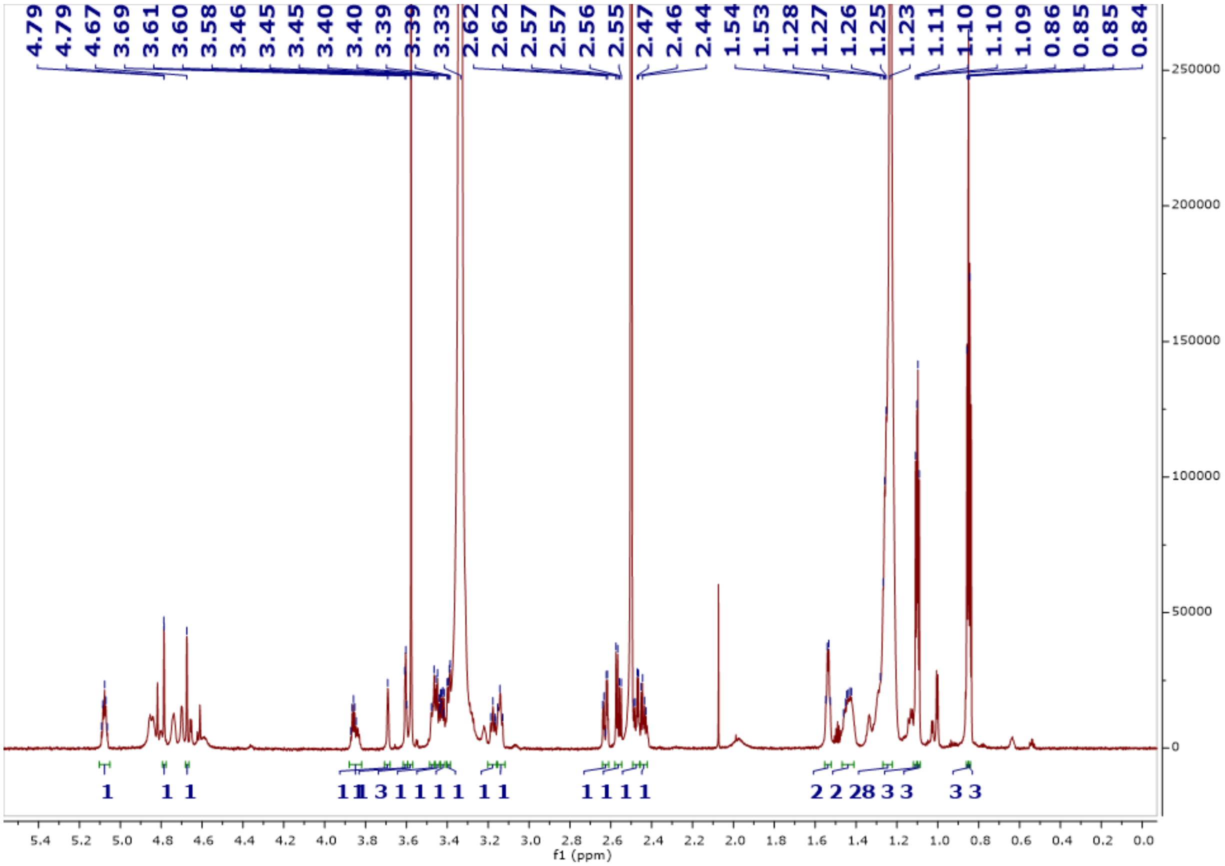

**Figure S1.** $^{1}H$ NMR spectrum of rhamnolipid methyl ester A, recorded in DMSO-$d_6$ at 900 MHz.

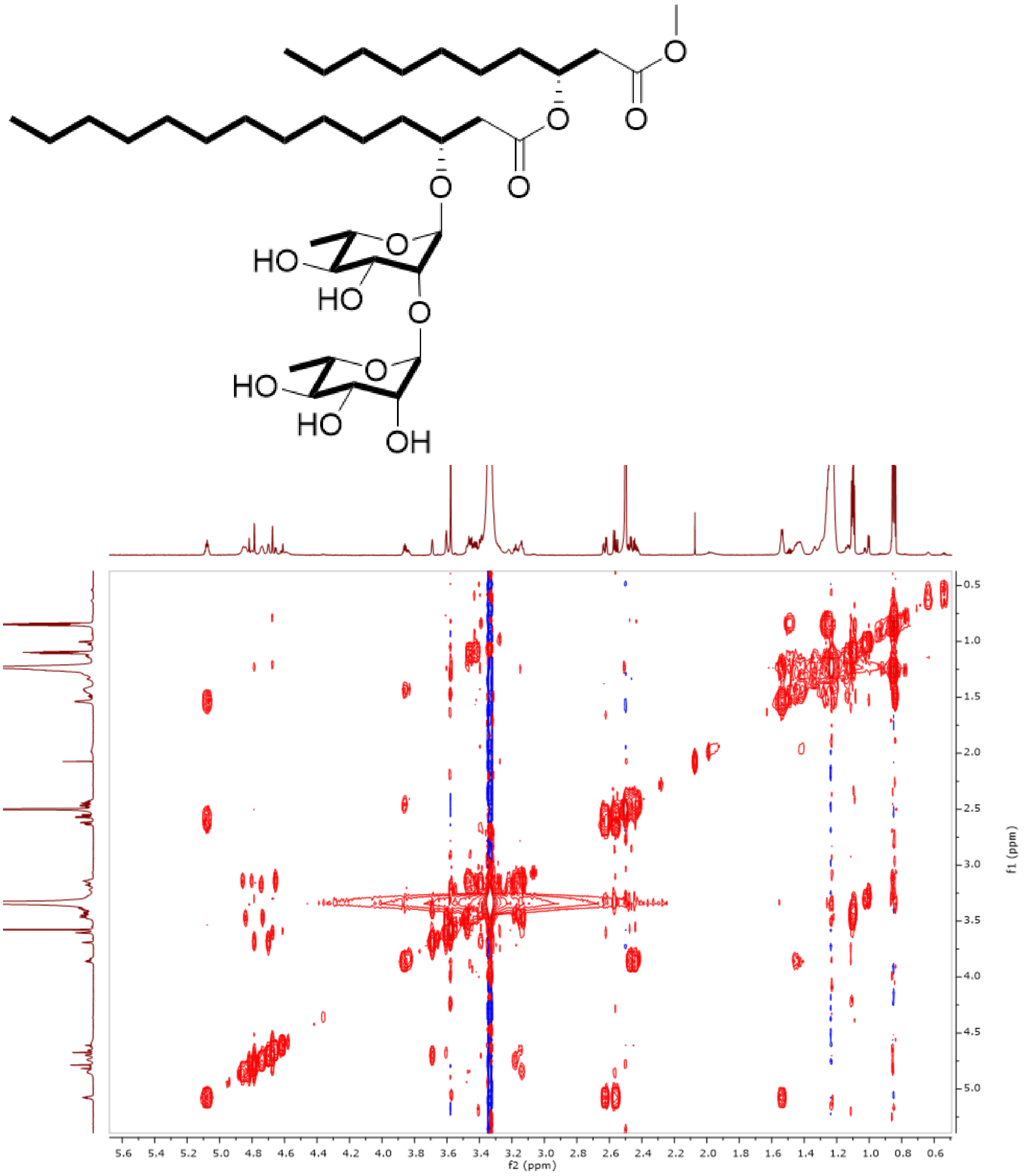


**Figure S2.** $^{1}H$-$^{1}H$ COSY spectrum of rhamnolipid methyl ester, recorded in DMSO-$d_6$ at 900 MHz.

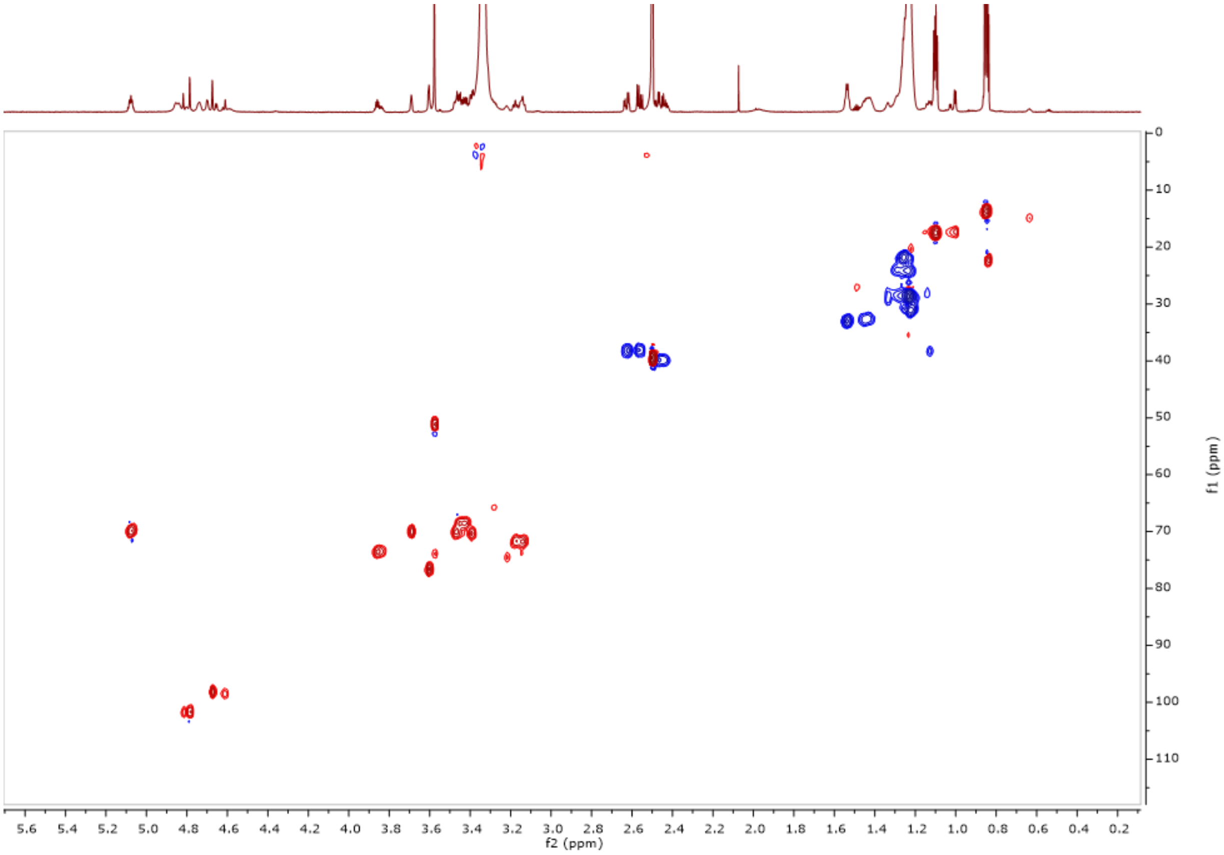


**Figure S3.** $^{1}H$-$^{13}C$ HSQC spectrum of rhamnolipid methyl ester, recorded in DMSO-$d_6$ at 900 MHz.

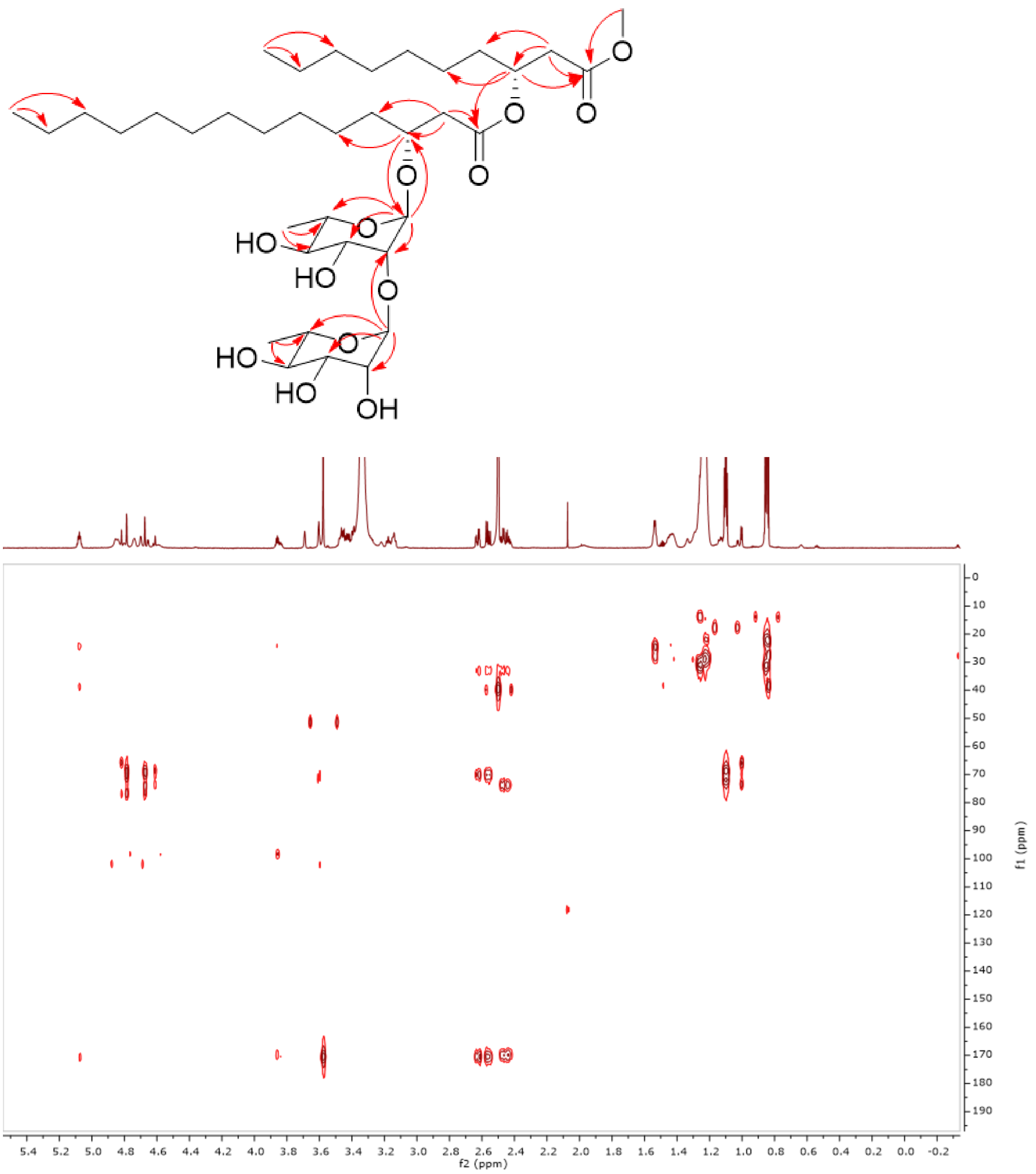


**Figure S4.** $^{1}H$-$^{13}C$ HMBC spectrum of rhamnolipid methyl ester, recorded in DMSO-$d_6$ at 900 MHz.

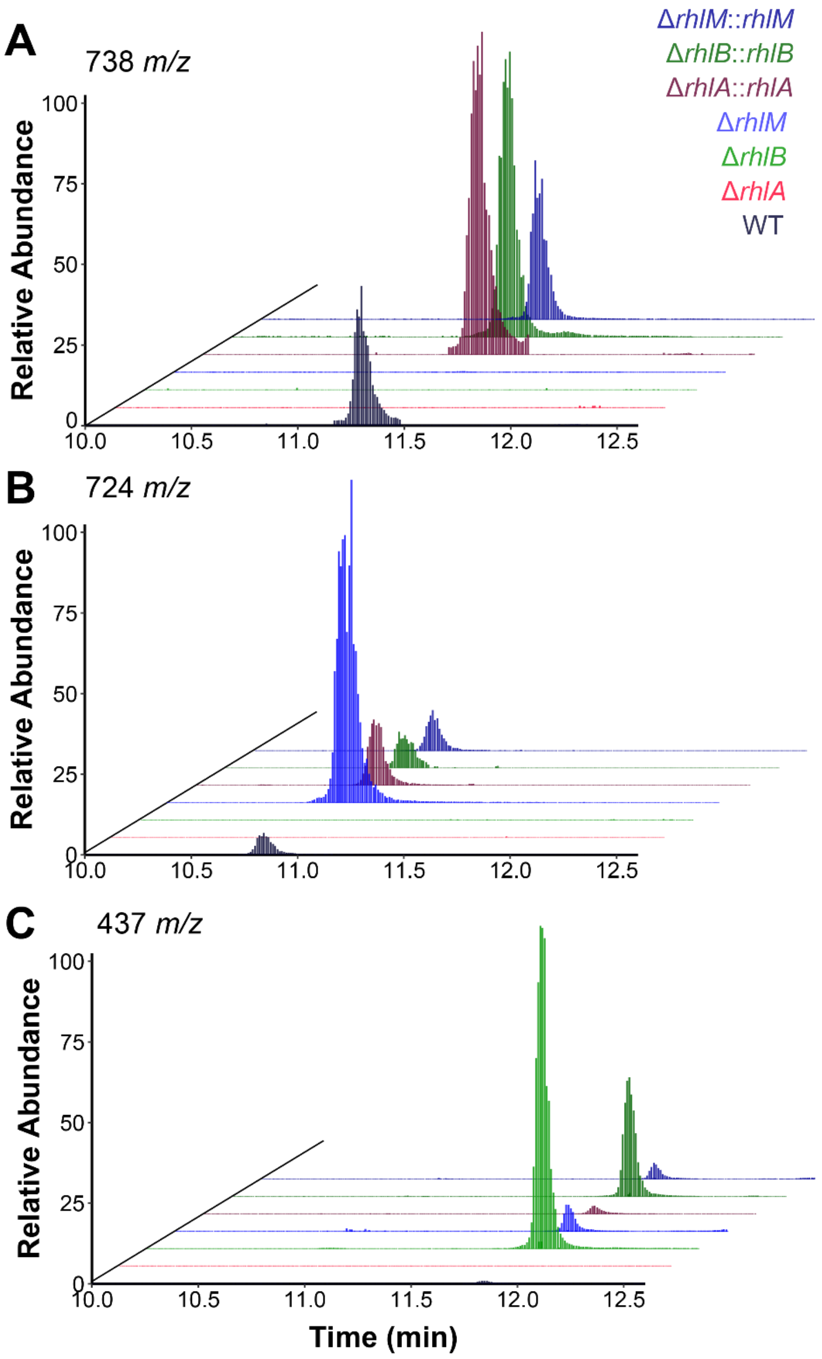


**Figure S5.** Extracted ion chromatograms (EIC) for *P. caledonica* F3 *rhl* mutants and genetic complement strains. (A) EIC for the RLME A ammonium adduct, 738 *m/z*, shows the absence of RLME A detected in three *rhl* knockout strains. RLME A production is rescued in genetic complementation strains. (B) EIC for the desmethyl RL ammonium adduct, 724 *m/z*, shows the accumulation of this intermediate in Δ*rhlM*, with low abundance detected in WT and complementation strains. (C) EIC for the HAA ammonium adduct, 437 *m/z*, shows the accumulation of this intermediate in Δ*rhlB*, with low abundance detected in WT and complementation strains. For each ion, intensities were normalized to the highest intensity detected among all strains.

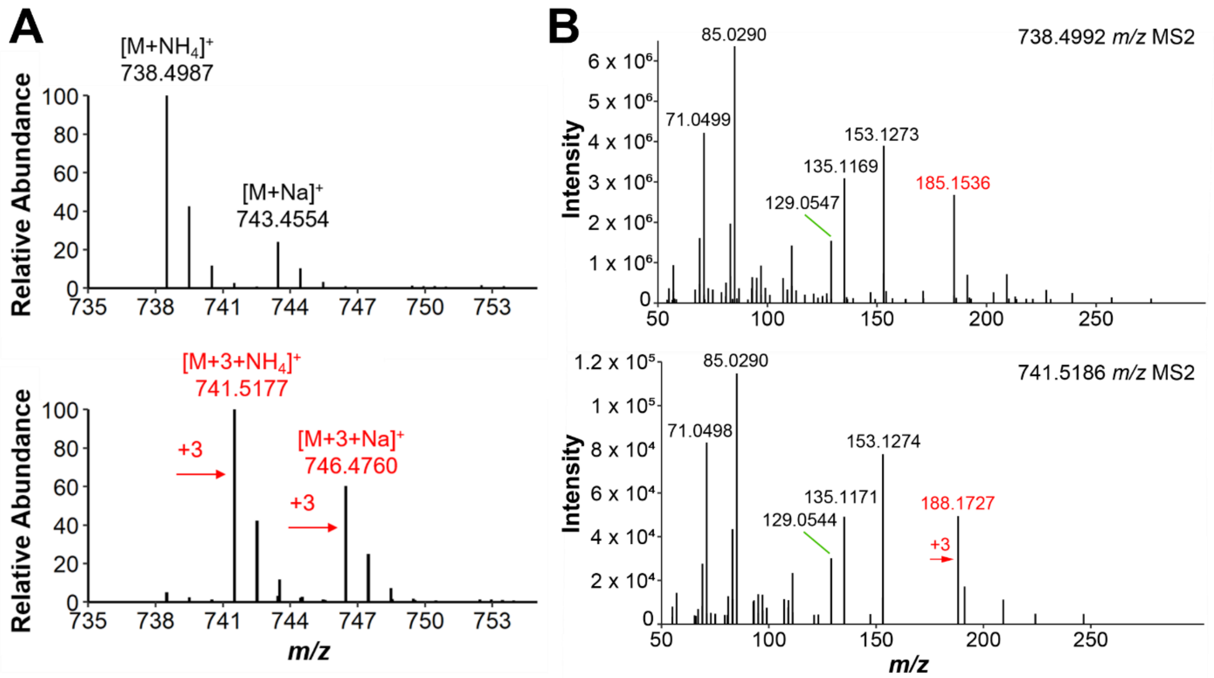


**Figure S6.** (A) Stable isotope labeling using $D_3$-Methionine verifies the biological origin of the carboxymethyl group, suggesting SAM-mediated methyltransferase activity. The +3 major isotopologues were observed for both ammoniated and sodiated adducts of RLME A in the labeled sample (bottom) compared to the unlabeled sample (top). (B) MS/MS spectra for the ammoniated adduct in the unlabeled sample (top) and labeled sample (bottom). Fragment 188.1537 *m/z*, corresponding to the acyl fragment containing the methyl ester, is the only fragment that shows incorporation of the $D_3$ label.

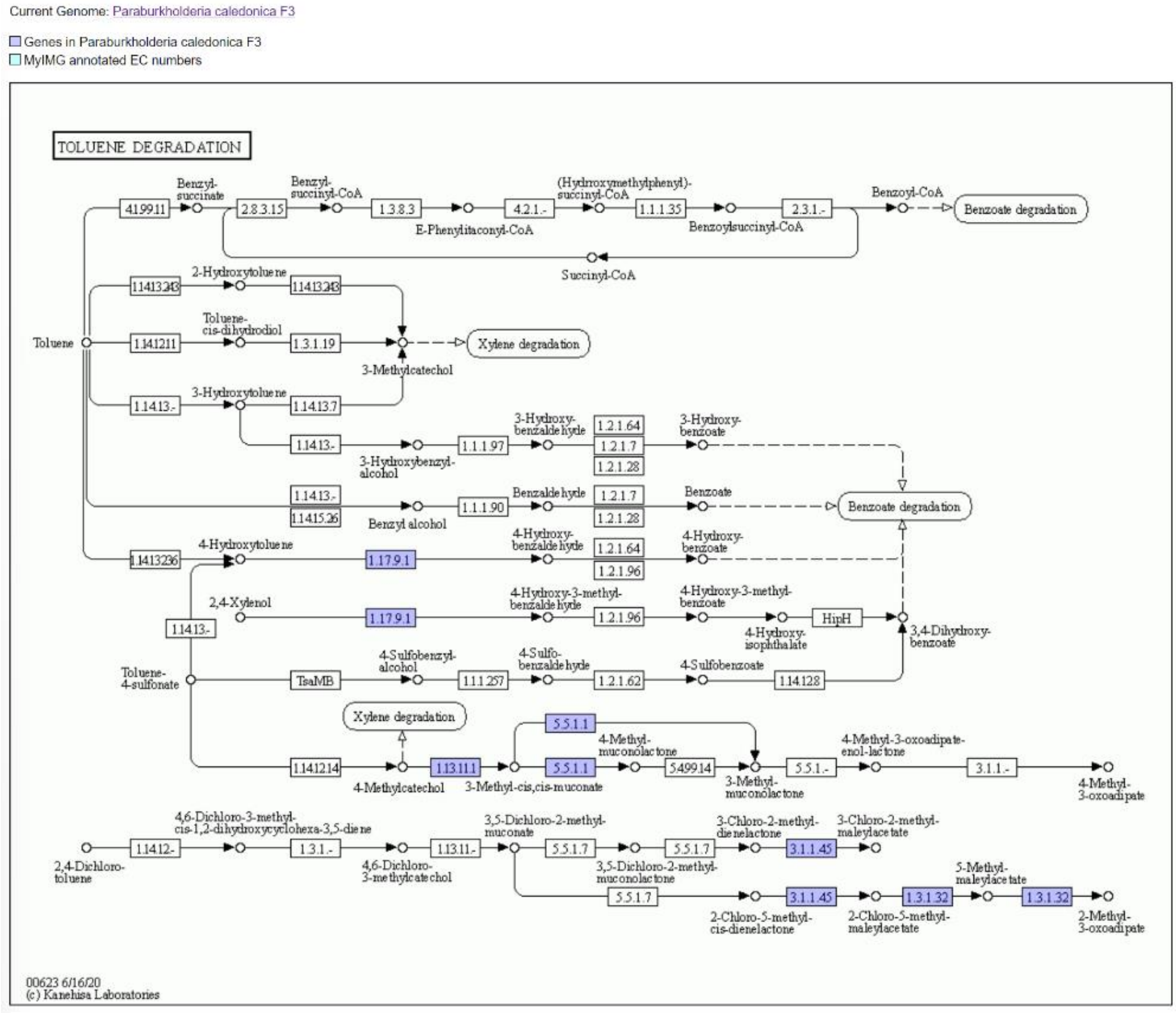


**Figure S7.** Predicted pathways for aromatic compound degradation in *P. caledonica* F3: toluene degradation pathways. KEGG analysis performed using JGI IMG-MER. Genes highlighted in lavender represent those found in the *P. caledonica* F3 genome.

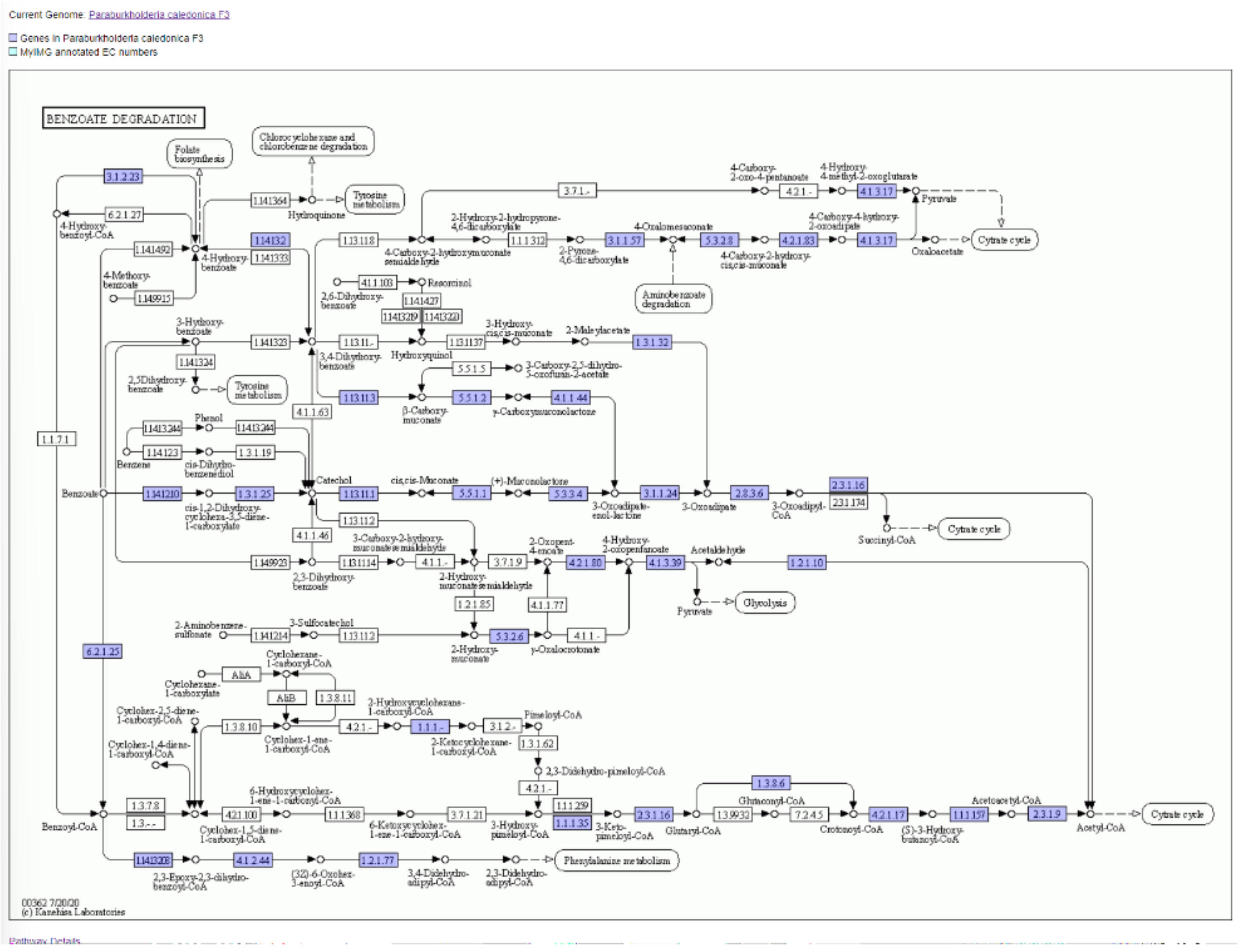


**Figure S8.** Putative pathways for aromatic compound degradation in *P. caledonica* F3: benzoate degradation pathways. KEGG analysis performed using JGI IMG-MER. Genes highlighted in lavender represent those found in the *P. caledonica* F3 genome.

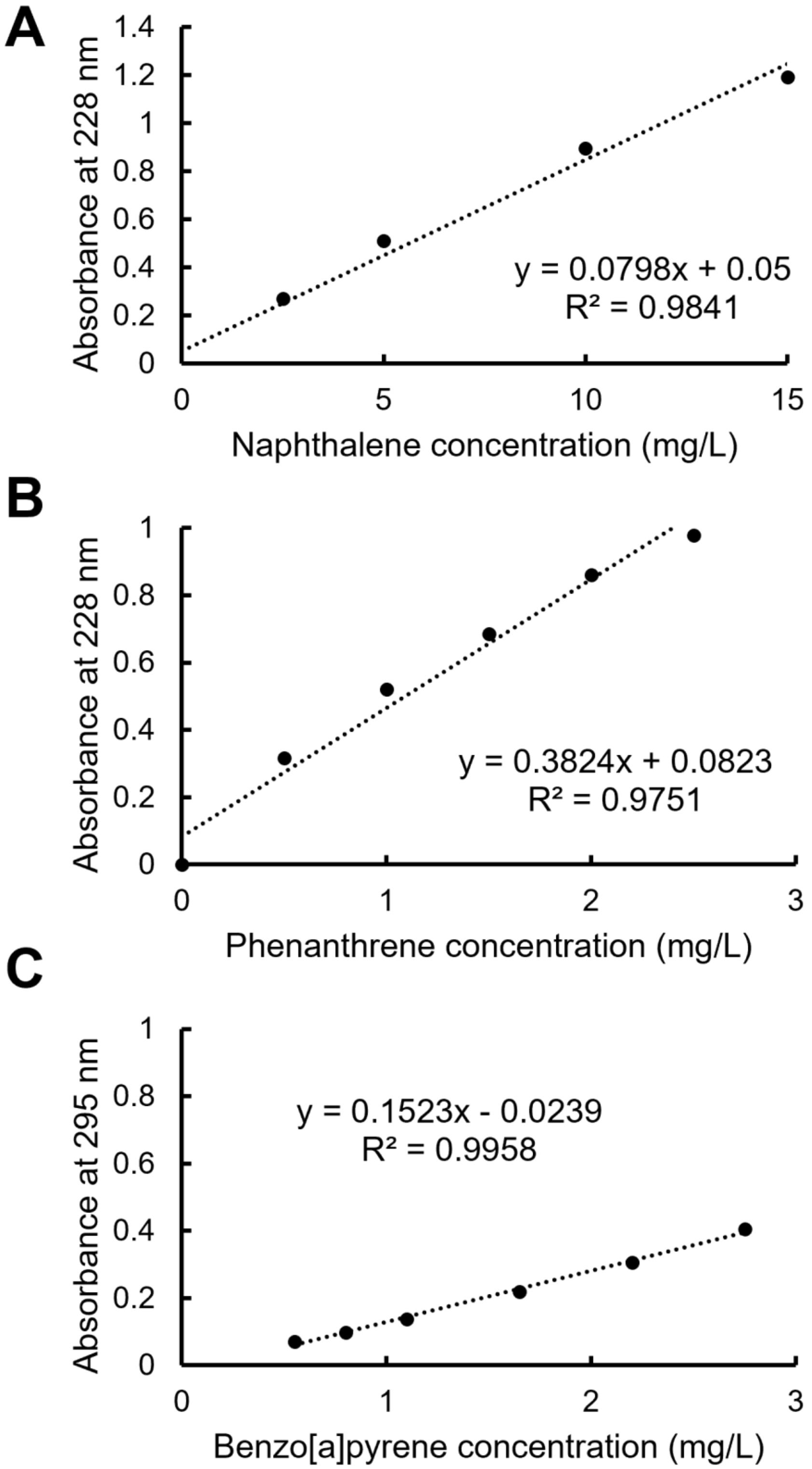


**Figure S9.** Standard curves for naphthalene (A), phenanthrene (B), and benzo[a]pyrene (C) used in PAH solubilization experiments.

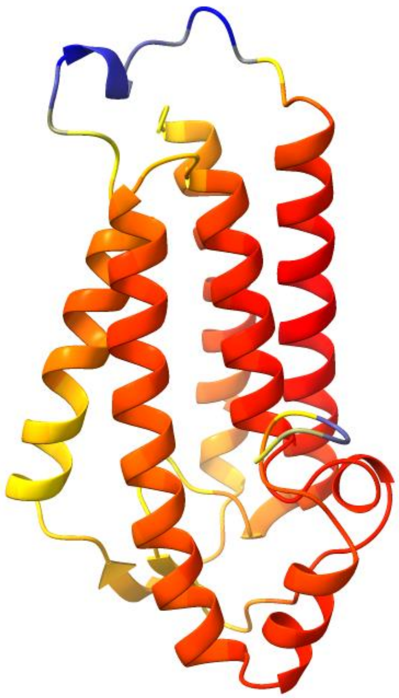

**Figure S10.** AlphaFold structure of RhlM colored by pLDDT score. Threshold values of 58.23, 77.75, and 98.11 were used for blue, yellow, and red, respectively.

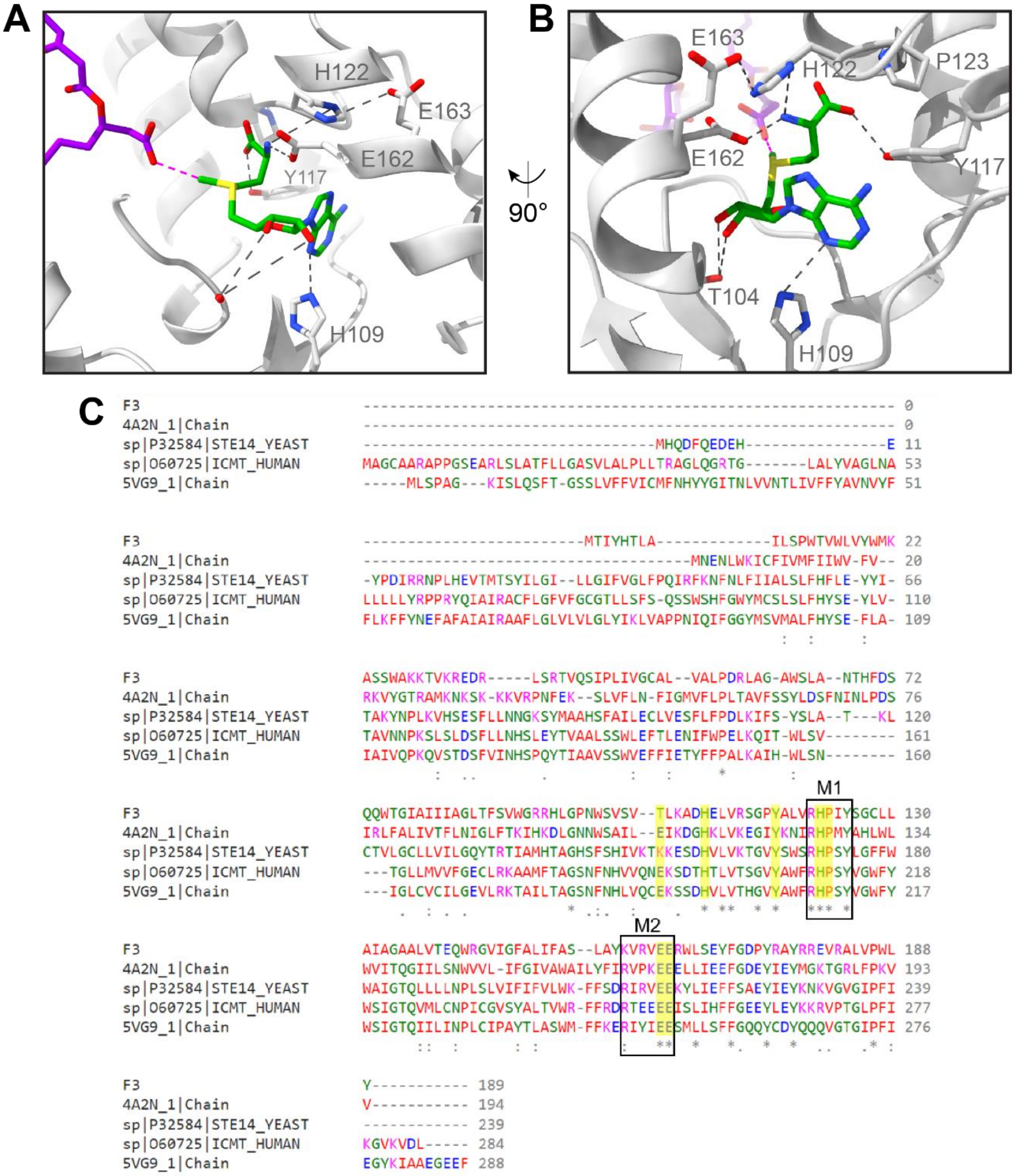


**Figure S11. (A, B)** Interactions with S-adenosyl methionine (SAM, green sticks). Putative interacting residues are shown as sticks. The cofactor is enclosed by conserved regions of RhlM. Gray dashed lines indicate hydrogen bonds. Magenta dashed line indicates measured distance between RL carboxylate O (methyl acceptor) and SAM methyl C (methyl donor). Atom coloring is: nitrogen, blue; oxygen, red; sulfur, yellow. (C) Multiple sequence alignment of RhlM (F3), Ma Mtase (4A2N), *Saccharomyces cerevisiae* ICMT (STE14_YEAST), *Homo Sapiens* ICMT (ICMT_HUMAN), and Tc ICMT (5VG9) made with Clustal Omega. M1 and M2 are conserved motifs within the cofactor-binding pocket of ICMT family proteins. Residues highlighted in yellow correspond to interacting residues shown in A and B.

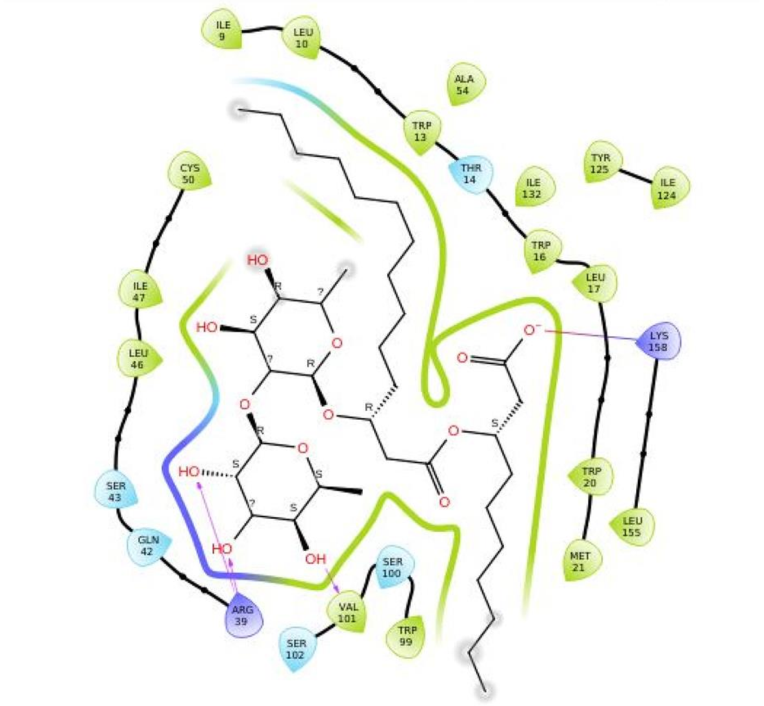


**Figure S12.** A 2D schematic ligand interaction diagram for RL and RhlM. Residues and the cartoon contact strip are colored by interaction type: green = hydrophobic, light blue = polar, dark blue = charged (positive). Atoms shaded with gray circles are solvent exposed.
